# Lrfn2-mutant mice display suppressed synaptic plasticity and inhibitory synapse development and abnormal social communication and startle response

**DOI:** 10.1101/252429

**Authors:** Yan Li, Ryunhee Kim, Yi Sul Cho, Doyoun Kim, Kyungdeok Kim, Junyeop Daniel Roh, Hanwool Park, Esther Yang, Soo-Jeong Kim, Jaewon Ko, Hyun Kim, Yong-Chul Bae, Eunjoon Kim

## Abstract

SALM1, also known as LRFN2, is a PSD-95-interacting synaptic adhesion molecule implicated in the regulation of NMDA receptor (NMDAR) clustering largely based on in vitro data, although its in vivo functions remain unclear. Here, we found that mice lacking SALM1/LRFN2 (*Lrfn2*^*-/-*^ mice) show a normal density of excitatory synapses but altered excitatory synaptic function, including enhanced NMDAR-dependent synaptic transmission but suppressed NMDAR-dependent synaptic plasticity in the hippocampal CA1 region. Unexpectedly, SALM1 expression is detected in both glutamatergic and GABAergic neurons, and *Lrfn2*^*-/-*^ CA1 pyramidal neurons show decreases in the density of inhibitory synapses and frequency of spontaneous inhibitory synaptic transmission. Behaviorally, ultrasonic vocalization was suppressed in *Lrfn2*^*-/-*^ pups separated from their mothers, and acoustic startle was enhanced, but locomotion, anxiety-like behavior, social interaction, repetitive behaviors, and learning and memory were largely normal in adult *Lrfn2*^*-/-*^ mice. These results suggest that SALM1/LRFN2 regulates excitatory synapse function, inhibitory synapse development, and social communication and startle behaviors in mice.

**Significance Statement:** Synaptic adhesion molecules regulate synapse development and function, which govern neural circuit and brain functions. The SALM/LRFN family of synaptic adhesion proteins consists of five known members whose in vivo functions are largely unknown. Here we characterized mice lacking SALM1/LRFN2 (SALM1 knockout) known to associate with NMDA receptors and found that these mice showed altered NMDA receptor-dependent synaptic transmission and plasticity, as expected, but unexpectedly also exhibited suppressed inhibitory synapse development and synaptic transmission. Behaviorally, SALM1 knockout pups showed suppressed ultrasonic vocalization upon separation from their mothers, and SALM1 knockout adults showed enhanced responses to loud acoustic stimuli. These results suggest that SALM1/LRFN2 regulates excitatory synapse function, inhibitory synapse development, social communication, and acoustic startle behavior.

## Introduction

Synaptic adhesion molecules with the prototypical molecules being neuroligins and neurexins have been shown to regulate the development, function, and plasticity of neuronal synapses (Dalva et al., 2007; Biederer and Stagi, 2008; Shen and Scheiffele, 2010; Siddiqui and Craig, 2011; Krueger et al., 2012; Missler et al., 2012; Valnegri et al., 2012; Takahashi and Craig, 2013; Um and Ko, 2013; Bemben et al., 2015; Ko et al., 2015; de Wit and Ghosh, 2016; Jang et al., 2017; Sudhof, 2017; Um and Ko, 2017; Yuzaki, 2017). Among such synaptic adhesion molecules is the leucine-rich repeat (LRR)-containing SALM/LRFN (synaptic adhesion-like molecule/leucine rich repeat and fibronectin type III domain containing) family comprising five known members: SALM1/LRFN2, SALM2/LRFN1, SALM3/LRFN4, SALM4/LRFN3, and SALM5/LRFN5 (Ko et al., 2006; Morimura et al., 2006; Wang et al., 2006; Nam et al., 2011).

SALM family proteins share a similar domain structure, consisting of six LRRs, an Ig domain, and a fibronectin type III domain in the extracellular region, followed by a single transmembrane domain and a cytoplasmic region. The extreme C-terminal tails of SALM1–3, but not SALM4/5, contain a PDZ domain-binding motif that interacts with the PDZ domains of PSD-95, an abundant postsynaptic scaffolding protein (Sheng and Kim, 2011; Won et al., 2017). The cytoplasmic regions of individual SALMs share minimal amino acid sequence identities, except for the C-terminal PDZ-binding motif, suggestive of functional diversity. However, SALMs associate with each other to form various homomeric and heteromeric complexes in vivo (Seabold et al., 2008; Lie et al., 2016), perhaps for concerted actions.

Functionally, SALMs regulate synapse development. SALM3 and SALM5, but not other SALMs, promote synapse development by interacting with presynaptic LAR family receptor protein tyrosine phosphatases (LAR-RPTPs) (Mah et al., 2010; Li et al., 2015; Choi et al., 2016). Mice lacking SALM3/LRFN4 display suppressed excitatory synapse development (Li et al., 2015). SALM4/LRFN3, despite lacking synaptogenic activity, interacts in cis with SALM3 and inhibits SALM3-dependent synapse development (Lie et al., 2016).

SALMs also regulate excitatory synaptic transmission and plasticity. SALM1 associates with NMDA receptors (NMDARs) and induces their dendritic clustering through a mechanism that requires its C-terminal PDZ-binding motif in vitro (Wang et al., 2006), but it does not interact with AMPA receptors (AMPARs). SALM2 promotes excitatory synapse maturation by associating with both NMDARs and AMPARs (Ko et al., 2006). Mice lacking SALM1/LRFN2 have recently been reported to display impairments in excitatory synapse maturation and enhancements in long-term potentiation (LTP) that are associated with autistic-like social deficits and stereotypies as well as enhanced learning and memory (Morimura et al., 2017).

SALMs have also been implicated in human brain disorders. SALM1/LRFN2 has been associated with autism spectrum disorders (ASDs) (Voineagu et al., 2011; Morimura et al., 2017), schizophrenia (Morimura et al., 2017), working memory deficits (Thevenon et al., 2016), and antisocial personality disorders (Rautiainen et al., 2016). However, how a SALM1/LRFN2 deficiency in humans leads to these abnormalities remains unclear.

In the present study, we further explored in vivo functions of SALM1 using an independent *Lrfn2*-knockout mouse line (*Lrfn2*^*–/–*^ mice) and found that SALM1 is important for excitatory synaptic plasticity, inhibitory synapse development, and ultrasonic vocalization (USV) and acoustic startle.

## Materials and methods

### cDNA constructs

Full-length human SALM1 (aa 1-788) in pcDNA3.1 Myc HisA vector has been described (Mah et al., 2010).

### Antibodies

SALM1 (2022) guinea pig polyclonal antibodies were generated using the last 30 amino acids of mouse SALM1 as immunogen (NGMLLPFEESDLVGARGTFGSSEWVMESTV). The NeuN antibody was purchased from Millipore.

### Generation and characterization of *Lrfn2*^*–/–*^mice

*Lrfn2*-deficient mice were generated by Biocytogen by targeting the exon 2 of the *Lrfn2* gene under the genetic background of C57BL/6J. To remove the EGFP + neo cassette, mice were crossed with *Protamine-Flp* mice. For global *Lrfn2* knockout in the whole body, mice removed of the EGFP + neo cassette were crossed with *Protamine*-Cre mice. The resulting mice were crossed with WT mice to produce heterozygous mice (*Lrfn2*^*+/–*^). Male and female *Lrfn2*^*+/–*^ mice were crossed to produce WT and *Lrfn2*^*–/–*^ mice for all the experiments performed except for X-gal experiments. Mice were PCR-genotyped using the following primers: for WT allele: 5’-ATGGAGACTCTGCTTGGTGGGC-3’ (forward) and 5’-GTTAGCAAGGAAGCCTGGGAGC-3’ (reverse); for KO allele: 5’- CCAAGTAACTAGGTTGTTCTGGGC-3’ and 5’-TGAGGAATCTGGAACCGACCAG-3’. The primers for RT-PCR were 5’- AATAAGCTGCTCAGGGCTCTC-3’ and 5’-CAGACAGATTCTGGCAGACG-3’. For X-gal staining, mouse sperms carrying the LacZ cassette in the *Lrfn2* gene in the genetic background of C57BL/6N Tac were purchased from KOMP (VG15208), and used to produce progenies for X-gal staining using in vitro fertilization with oocytes in the C57BL/6J Tac background. Heterozygous male mice were used for X-gal staining.

## X-gal staining

Mice (P49) were perfused transcardially with heparinized 1 x phosphate buffered saline (PBS) and 4% paraformaldehyde. Vibratome brain sections (250 μm thickness) were incubated in staining solution (5 mM K3Fe(CN)6, 5 mM K4Fe(CN)6•3H2O, 2 mM MgCl2, 0.01% deoxycholate,1 mg/mL X-gal, 0.02% NP-40 in PBS) for 3 hours at room temperature.

## Electron microscopy

WT and *Lrfn2*^*–/–*^ mice were deeply anesthetized with sodium pentobarbital (80 mg/kg, intraperitoneal) and were intracardially perfused with 10 ml of heparinized normal saline, followed by 50 ml of a freshly prepared fixative of 2.5% glutaraldehyde and 1% paraformaldehyde in 0.1 M phosphate buffer (PB, pH 7.4). Hippocampus was removed from the whole brain, postfixed in the same fixative for 2 hours and stored in PB overnight at 4 °C. Sections were cut transversely on a Vibratome at 70 μm. The sections were osmicated with 0.5% osmium tetroxide (in PB) for 1 hour, dehydrated in graded alcohols, flat embedded in Durcupan ACM (Fluka), and cured for 48 hours at 60°C. Small pieces containing stratum radiatum of hippocampal CA1 region were cut out of the wafers and glued onto the plastic block by cyanoacrylate. Ultrathin sections were cut and mounted on Formvar-coated single slot grids. For quantification of excitatory synapse sections were stained with uranyl acetate and lead citrate, and examined with an electron microscope (Hitachi H-7500; Hitachi) at 80 kV accelerating voltage. For quantification of inhibitory synapse, sections were further immunogold stained for GABA.

## Postembedding immunogold staining for GABA

Sections were immunostained for GABA by postembedding immunogold method, as previously described (Paik et al., 2007), with some modifications. In brief, the grids were treated for 5 min in 1% periodic acid, to etch the resin, and for 8 min in 9% sodium periodate, to remove the osmium tetroxide, then washed in distilled water, transferred to Tris-buffered saline containing 0.1% Triton X-100 (TBST; pH 7.4) for 10 min, and incubated in 2% human serum albumin (HSA) in TBST for 10 min. The grids were then incubated with rabbit antiserum against GABA (GABA 990, 1:10,000) in TBST containing 2% HSA for 2 hours at room temperature. The antiserum (a kind gift from professor O. P. Ottersen at the Center for Molecular Biology and Neuroscience, University of Oslo) was raised against GABA conjugated to bovine serum albumin with glutaraldehyde and formaldehyde (Kolston et al., 1992) and characterized by spot testing (Ottersen and Storm-Mathisen, 1984). To eliminate cross-reactivity, the diluted antiserum was preadsorbed overnight with glutaraldehyde (G)-conjugated glutamate (500 μM, prepared according as described previously) (Ottersen et al., 1986). After extensive rinsing in TBST, grids were incubated for 3 hours in goat anti-rabbit IgG coupled to 15 nm gold particles (1:25 in TBST containing 0.05% polyethylene glycol; BioCell Co., Cardiff, United Kingdom). After a rinse in distilled water, the grids were counterstained with uranyl acetate and lead citrate, and examined with an electron microscope (Hitachi H-7500; Hitachi) at 80 kV accelerating voltage. To assess the immunoreactivity for GABA, gold particle density (number of gold particles per μm^2^) of each GABA+ terminal was compared with gold particle density of terminals which contain round vesicles and make asymmetric synaptic contact with dendritic spines (background density). Terminals were considered GABA-immunopositive (+) if the gold particle density over the vesicle-containing areas was at least five times higher than background density.

## Quantitative analysis of excitatory and inhibitory synapses

For quantification of excitatory synapses, twenty-four electron micrographs representing 368.9 μm^2^ neuropil regions in each mouse were taken at a 40,000×. Number of spines (PSD density), proportion of perforated spines, PSD length, and PSD thickness from each of three WT and *Lrfn2*^*–/–*^ mice were quantified by using ImageJ software. For quantification of inhibitory synapses, twenty-four electron micrographs representing 655.5 μm^2^ neuropil regions in each mouse were taken at a 30,000×. Number of GABA+ terminals showing clear PSD (inhibitory synapse density), length and thickness of PSD contacting GABA+ terminals from each of three WT and *Lrfn2*^*–/–*^ mice were quantified by using ImageJ software. The measurements were all performed by an experimenter blind to the genotype. Digital images were captured with GATAN DigitalMicrograph software driving a CCD camera (SC1000 Orius; Gatan) and saved as TIFF files. Brightness and contrast of the images were adjusted using Adobe Photoshop 7.0 (Adobe Systems).

## In situ hybridization

Mouse brains at various developmental stages (embryonic day 18 and postnatal day 0, 7, 14, 21, and 42) were extracted and rapidly frozen in isopentane prechilled with dry ice. Brain sections were prepared with a cryostat and thaw-mounted onto gelatin-coated slides and fixed in 4% paraformaldehyde. Two independent hybridization probes targeted roughly the N‐ and C-term regions of the *Lrfn2* gene (NM_027452.3). For probe 1, the forward sequence was TACGCCGGATCCGTGGGCTGCTGGCTTTT and reverse sequence is TACGCCGAATTCTGGCTGATGGTGTTCCTG. For probe 2, the forward sequence is TACGCCGGATCCTGCTCTTGCCCTTTGAGG and reverse sequence is TACGCCGAATTCATGGGGAAGGGGGTGTAG.

## Fluorescent in situ hybridization

In brief, frozen sections (14 μm thick) were cut coronally through the hippocampal formation. Sections were thaw-mounted onto Superfrost Plus Microscope Slides (Fisher Scientific 12-550-15). The sections were fixed in 4% formaldehyde for 10 min, dehydrated in increasing concentrations of ethanol for 5 min, and finally air‐dried. Tissues were then pretreated for protease digestion for 10 min at room temperature. For RNA detection, incubations with different amplifier solutions were performed in a HybEZ hybridization oven (Advanced Cell Diagnostics) at 40 °C. In this study, we used fluorescent probes to label Lrfn2, Gad1/2 and Vglut1/2; mixtures of Gad1 + Gad2 probes, and Vglut1 + Vglut2 probes, were used to collectively label GABAergic and glutamatergic neurons, respectively. Synthetic oligonucleotides in the probes were complementary to the following nucleotide regions in the target genes; Lrfn2, nucleotide sequence 542 – 2041 of Mm-Lrfn2-C1; Vglut1, 464 – 1415 of Mm‐ Slc17a7-C2, Vglut2, 1986 – 2998 of Mm-Slc17a6-C2, Gad1, 62 – 3113 of Mm‐ Gad1-C3, Gad2, 552 – 1506 of Mm-Gad2-C3, (Advanced Cell Diagnostics), respectively. The labeled probes were conjugated to Atto 550 (C1), Atto 647 (C2), and Alexa Fluor 488 (C3). The sections were hybridized at 40 °C with labeled probe mixtures (C1 + C2 + C3) per slide for 2 hours. Then the nonspecifically hybridized probes were removed by washing the sections, three times each in 1x wash buffer at room temperature for 2 min. Amplification steps involved sequential incubations with Amplifier 1-FL for 30 min, Amplifier 2-FL for 15 min, Amplifier 3-FL for 30 min, and Amplifier 4 Alt B-FL at 40 ^o^C for 15 min. Each amplifier solutions were removed by washing three times with 1x wash buffer for 2 min at RT. Fluorescent images were acquired using LSM 700 microscope (Zeiss) and analyzed using ImageJ software.

## Electrophysiology

For whole-cell patch-clamp analysis, sagittal hippocampal and coronal mPFC slices (300 μm thick) from *Lrfn2*^*–/–*^ mice and their wild-type (WT) littermates at 21–24 postnatal days were prepared using a vibratome in ice-cold section buffer containing (in mM) 212 sucrose, 25 NaHCO_3_, 5 KCl, 1.25 NaH_2_PO_4_, 0.5 CaCl_2_, 3.5 MgSO_4_, 10 D-glucose, 1.2 L-ascorbic acid, and 2 Na-pyruvate bubbled with 95% O_2_/5% CO_2_. The slices were recovered at 32 °C for 30 min in normal artificial CSF (ACSF) (in mM: 124 NaCl, 2.5 kCl, 1NaH_2_PO_4_, 25 NaHCO_3_, 10 glucose, 2 CaCl_2_, 2 MgSO_4_ oxygenated with 95% O_2_/5% CO_2_). Stimulation and recording pipettes were pulled from borosilicate glass capillaries (Harvard Apparatus) using a micropipette electrode puller (Narishege). Whole-cell patch-clamp recordings were made using a MultiClamp 700B amplifier (Molecular Devices) and Digidata 1440A (Molecular Devices) under visual control with differential interference contrast illumination in an upright microscope. Signals were filtered at 2 kHz and digitized at 10 kHz. Miniature EPSCs (mEPSCs) were obtained at a holding potential of -70 mV using patch electrode (3–4 MΩ) filled with internal solution (in mM): 100 CsMeSO_4_, 10 TEA-Cl, 8 NaCl, 10 HEPES, 5 QX-314-Cl, 2 Mg-ATP, 0.3 Na-GTP, 10 EGTA, with pH 7.25, 295 mOsm. TTX (0.5 μM) and picrotoxin (100 μM) were added to ACSF to inhibit spontaneous action potential-mediated synaptic currents and IPSCs, respectively. For spontaneous EPSCs (sEPSCs), the same experimental conditions used for mEPSC measurements were used, except for omitting TTX. To measure miniature IPSCs (mIPSCs), cells were also held at -70 mV, and pipette internal solution contained (in mM): 115 CsCl, 10 TEA-Cl, 8 NaCl, 10 HEPES, 5 QX-314-Cl, 4 Mg-ATP, 0.3 Na-GTP, 10 EGTA, with pH 7.35, 295 mOsm. To inhibit excitatory synaptic currents, TTX (0.5 μM), D-AP5 (25 μM), and NBQX (10 μM) were added to ACSF. sIPSC measurements were made in the absence of TTX.

For measurements of the NMDA/AMPA ratio, sagittal hippocampal slices (300 μm thick) were prepared. The recording pipettes (3–4 MΩ) were filled with the same internal solution used for mEPSC measurements. Picotoxin (100 μM) were added to ACSF. CA1 pyramidal neurons were voltage clamped at -70 mV, and EPSCs were evoked at every 15 sec. AMPAR-mediated EPSCs were recorded at -70 mV, and 20 consecutive responses were recorded after stable baseline. After recording AMPA receptor-mediated EPSCs, holding potential was changed to +40 mV to record NMDA receptor-mediated EPSCs. NMDA component was measured at 60 ms after the stimulation. The NMDA/AMPA ratio was calculated by dividing the mean value of 20 NMDA-EPSC peak amplitudes by the mean value of 20 AMPAEPSC peak amplitudes. Data were acquired using Clampex 10.4 (Molecular Devices) and analyzed using Clampfit 10.4 (NMDA/AMPA ratio; Molecular Devices).

For field recordings, sagittal hippocampal slices (400 mm thick) were prepared. The pipettes were filled with ACSF. For input/output and paired pulse ratio experiments, CA1 field excitatory postsynaptic potential (fEPSP) was evoked every 20 s and the stable baseline was recorded for 10 min. For Input/output recording, gradually increasing stimuli were delivered to induce fiber volley amplitudes of 0.05– 0.3 mV/ms. For paired pulse ratio recording, inter-stimulus intervals were 25, 50, 100, 200 and 300 ms. For LTP measurements, the Schaffer collateral pathway was stimulated every 20 s and a stable baseline was maintained for 20 min. LTP stimuli were high-frequency stimulation (100 Hz, 1s) or theta-burst stimulation (10 trains of 4 pulses at 100 Hz, delivered at 5 Hz, repeated 4 times at 10s interval). For LTD experiments, picrotoxin (100 μM) were added to ACSF, and low-frequency stimulation (1 Hz, 900 pulses) was given. After LTP or LTD stimulus, the responses were recorded for an hour.

## Animal behavioral tests

Male mice at 2∼6 months of age were used for all behavioral tests, except for pup retrieval test. All mice were fed ad libitum and housed under 12 h light/dark cycle, and all mouse behaviors were performed using mice at their light-off/dark periods. All procedures were approved by the Committee of Animal Research at KAIST(KA2012-19). Animal used in behavioral test were generated from crossing male and female heterozygous mice. All tests used littermates or age-matched mice.

## Open field test

Mice were placed into a white 40 x 40 x 40 cm open field box. Mice were allowed to explore freely in the box for an hour under complete darkness (∼0 lux) or 110 lux condition. Mouse movements were recorded using a video camera (infra-red camera in the case of complete darkness) and analyzed using the EthoVision XT 10 software (Noldus).

## Laboras^TM^ monitoring of movements

Home cage locomotion behaviors of mice were recorded and analyzed using Laboratory Animal Behavior Observation Registration and Analysis System (LABORAS^TM^, Metris) for 72 consecutive hours.

## Elevated plus-maze

The maze consisted of 2 open arms, 2 closed arms, and a center zone, elevated to a height of 50 cm above the floor. Mice were initially placed on the center zone faced to the open arm and allowed to freely explore the space for 8 min. Light condition was about 80 lux.

## Light-dark test

The apparatus had a dimension of 12 x 30 x 20 cm for the light chamber (∼600 lux) and 14 x 13 x 20 cm for the dark chamber (∼5 lux). Mice were placed in the center of the light chamber and allowed to explore the whole apparatus freely for 10 min. Time spent in the light chamber was analyzed using EthoVision XT 10.

## Three chamber test

Subject mice were isolated for 3 days in their home cages before the experiment. The apparatus consisted of 3 chambers, and both side chambers had a steel wire cage in the corner to place inanimate objects or mice. First, mice were put in the center zone and allowed to freely explore the whole apparatus for 10 min. Next, a stranger mouse(S1) was placed in a wire cage in a side chamber, and an inanimate object (O) was placed in another wire cage. Mice were then allowed to explore freely for 10 min. Stranger was randomly positioned in the left or right chamber. Then, the object was replaced with a novel stranger mouse (S2), and mice were allowed to freely explore the apparatus for 10 min. Time spent in each chamber and time spent sniffing the wire cage containing either O, S1 or S2 were analyzed using EthoVision XT 10.

## Direct interaction test

Subject mice were isolated for 4 days in their home cages. On day 1 for habituation, mice were individually placed in a grey box (30 cm x 30 cm x 40 cm) for 10 min. Twenty four hours later, two age-matched mice in the same genotype that never encountered each other before were put into the habituated box and allowed to interact with each other freely for 10 min. Physical interaction, nose-to-nose sniffing, following behaviors were measured manually in a blind manner.

## Ultrasonic vocalization (USV) test

For pup USVs, pups were placed in a glass bowl in a recording chamber and a recording microphone was placed 20 cm above the pup. USVs from pups induced by separation from their mothers were recorded using Avisoft Ultrasoundgate (Model 116Hb) system for 3 min. For adult USVs, male mice isolated for three days in their home cages, and these cages with mice were placed in a chamber with a microphone 20 cm above the home cage. Age-matched female mice were introduced to the home cage for 5 min. Recorded sound files were analyzed using the Avisoft SASLab Pro software (Avisoft Bioacoustics).

## Pup retrieval assay

Virgin female mice at 4 months of age were isolated in their home cages with nesting block for 4 days before the test. Three WT P1 pups were placed at three different corners away from the nesting block of the home cage of the test female mouse, and the female mouse was allowed to retrieve the pups for 10 min. Latency to each pup retrieval to the nest was measured.

## Repetitive behavior

Self-grooming and digging tests were performed in mouse home cages with fresh bedding. Mice were individually placed into a home cage for 20 min, and repetitive behaviors during the last 10 min were used to analyze self-grooming and digging behavior manually.

## Marble burying test

Subjected mice were placed in a home cage with 5-cm-thick beddings and 21 metal marbles and allowed to explore freely for 30 min. The analysis counted the number of buried marbles in a blind manner with marble burying counted when more than the two-thirds of marbles are buried.

## Rotarod test

Rotating speed of rod was gradually increased from 4 to 40 rpm over 5 min. Mice were placed gently on the rotating rod for 20 sec, followed by the start of rod rotation. The experiment was performed for 5 consecutive days, while measuring the latencies of mice to fall from the rod.

## Object recognition test

This test was performed in an open field test apparatus. Mice were habituated in the apparatus for 1 hour a day before training. For the displaced object recognition test (DORT), after exploring two same objects for 10 min, mice were put back to home cages for 5 min. One of the two objects was translocated to a position opposite in the box. Then, the mice were placed back in the apparatus and allowed to explore the objects for 10 min. Exploration time for the translocated object was measured. For the novel object test (NORT), on the training day, mice were allowed to explore two same objects for 10 min. 24 hours after training, one of the two objects was replaced with a new one, and mice were allowed to explore both objects freely for 10 min. Object exploration was defined by the amount of time spent for each object, with the nose of mice touched or faced towards the objects within 2 cm from them.

## Morris water maze

This assay was performed in a stainless round tank (12 cm diameter) with a hidden platform. The tank was filled with tap water at temperature of 20∼22 °C made opaque with white watercolor. For memory acquisition, mice were trained to find the platform with 3 trials per day with an inter-trial interval of 30 min for 5 days. When mice reached the platform, they were allowed to rest on the platform for 15 secs before they are put back to their home cages. If mice did not find the platform within 60 secs, they were guided to the platform and allowed to rest on the platform for 15 secs. On day 6, for probe test, the platform was removed and mice were put in the center of the tank and allowed to explore for 1 min. Mice were retrained to find the hidden platform in the tank after the probe test to avoid memory extinction. On the next day (day 7), the platform was replaced to a site opposite to the original position, and mice were trained for 4 days for reversal learning and memory. On day 11, another probe test was performed to test reversal learning and memory. The number of exact crossing over the platform region, quadrant occupancy, and swimming speed were analyzed using the EthoVision XT 10.

## Fear conditioning

On the training day, mice were placed in the fear chamber and allowed to explore the chamber freely for 2 min. Then, the mice went through five rounds of a 20 sec tone with a 0.5-mA foot shock during the last two secs followed by 40 sec rest. The final shock was followed by a 2-min post-training habituation. 24 hours later, mice were returned to the same shock chamber for 5 min to test contextual fear conditioning. Four hours later, the mice were returned to the chamber with a different context to test cued fear conditioning. To change the context, mice were placed in a round-shape tube added to the chamber where mice were allowed to explore freely for 2 min. Then, a 3-min tone was given to test levels of cued fear conditioning.

## Acoustic startle and pre-pulse inhibition

Different mouse cohorts were used for acoustic startle responses and pre-pulse inhibition (Wells et al., 2016). To test acoustic startle responses, the session was preceded by a 5-min exposure to 65 dB background noise. Then each mouse received 92 trials with inter-trial intervals ranging from 7-23 sec in pseudo-random order. The trials included a presentation of eight pulse-alone trials (120 dB, 40 ms pulse, four were given at the beginning and four at the end of the test), 77 pulse trials (seven each of 70, 75, 80, 85, 85, 90, 95,100, 105, 110, 115, and 120 dB, 40 ms pulse), and seven trials each without pulse or pre-pulse inhibition. To test pre-pulse inhibition, each mouse received 57 trials with inter-trial intervals ranging from 7-23 sec presented in pseudo-random order. The trials included a presentation of eight pulse-alone trials (120 dB, 40 ms pulse, four were given at the beginning and four at the end of the test), 35 pre-pulse trials (seven each of 70, 75, 80, 85 and 90 dB, 20 ms pre-pulse given 100 ms before a 120 dB, 40 ms pulse), and seven trials each without pulse or pre-pulse presentation. The pre-pulse inhibition percentage was calculated as follows: (100-(mean pre-pulse response/mean pulse response) x 100)). Startle at each pulse level was averaged across trials.

## PTZ-induced seizure

Mice were injected of 50 μg/weight (g) pentylenetetrazole (PTZ) into the intraperitoneal cavity and recorded of seizure for 10 min. Seizures were scored blindly according to Racine scale designed for PTZ-induced seizures in mice (Ferraro et al., 1999).

## Experimental design and statistical analysis

The order of behavioral tests was designed in a way to minimize stress in animals. The behavioral experiments were performed in the following order using three independent cohorts: cohort 1 underwent elevated plus maze, open field test (110 lux), digging and grooming, light-dark test, three chamber test, direct interaction test, PPI and fear conditioning test; cohort 2 underwent open field test (0 lux), adult USV, acoustic startle response, and PTZ-induced seizure; cohort 3 underwent Laboras test, displaced object recognition test, novel object recognition test, and Morris water maze. All statistical tests were performed using GraphPad Prism 7.0. Normality of data distribution was assessed using D’Agostino-Pearson omnibus test. Comparison of WT and KO data were performed using unpaired two-tailed Student’s t-test when the data showed Gaussian distribution, while Mann-Whitney test or Wilcoxon test was used when the data followed non-Gaussian distribution. Repeated measures two-way ANOVA was used to determine between-subject variable (genotype) and within-subject variable (repeated measures) for the analysis of the measures of input-output ratio, paired pulse facilitation, open field, Laboras, rotarod, Morris water maze, fear conditioning, acoustic startle, and paired pulse inhibition tests. Bonferroni’s test followed by ANOVA was used as a posthoc test for multiple comparisons. All data were displayed as mean ± SEM. The age, sex, and numbers of animals, and all the details of statistical results are shown in **Table 1**.

**Table 1.**
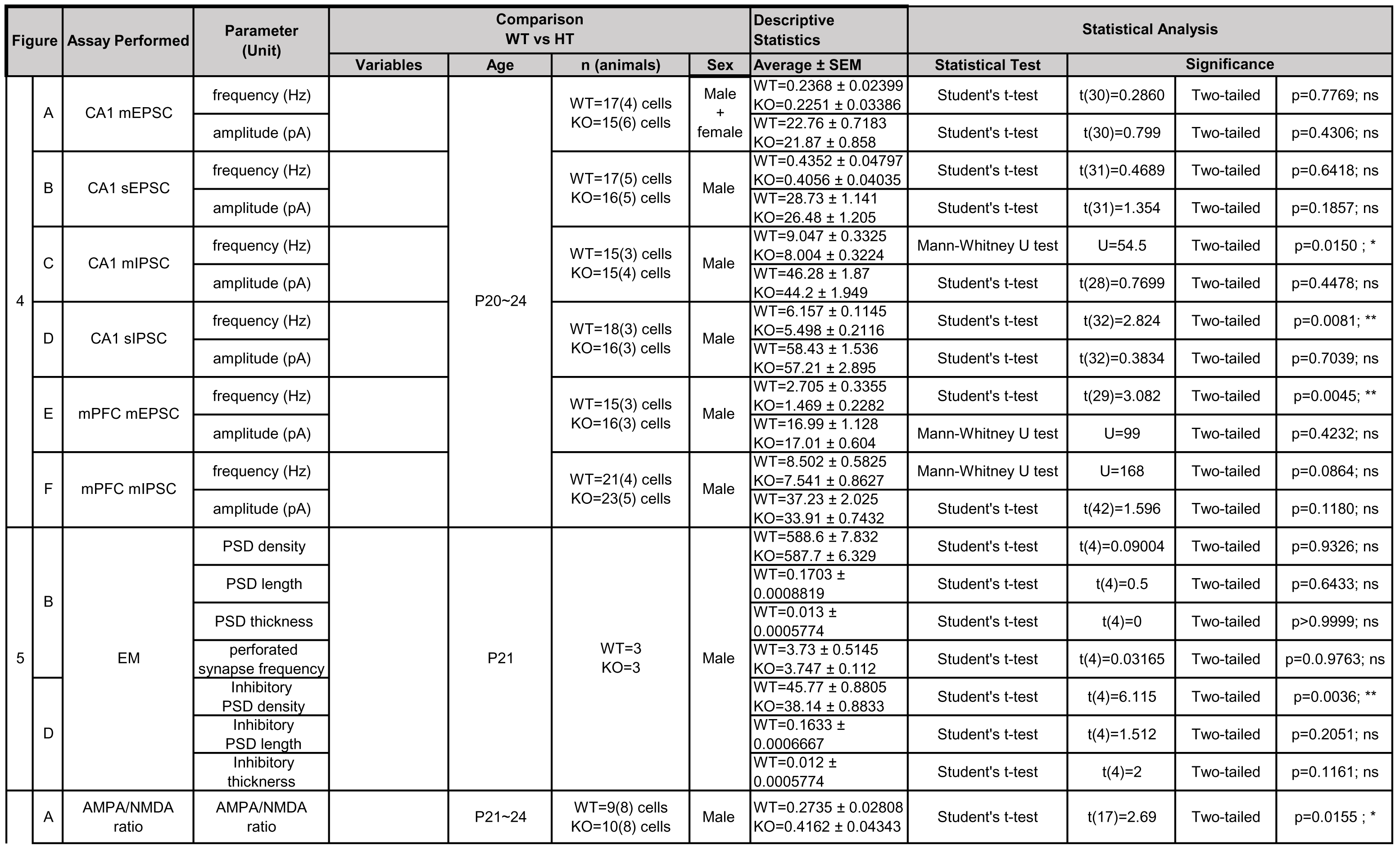
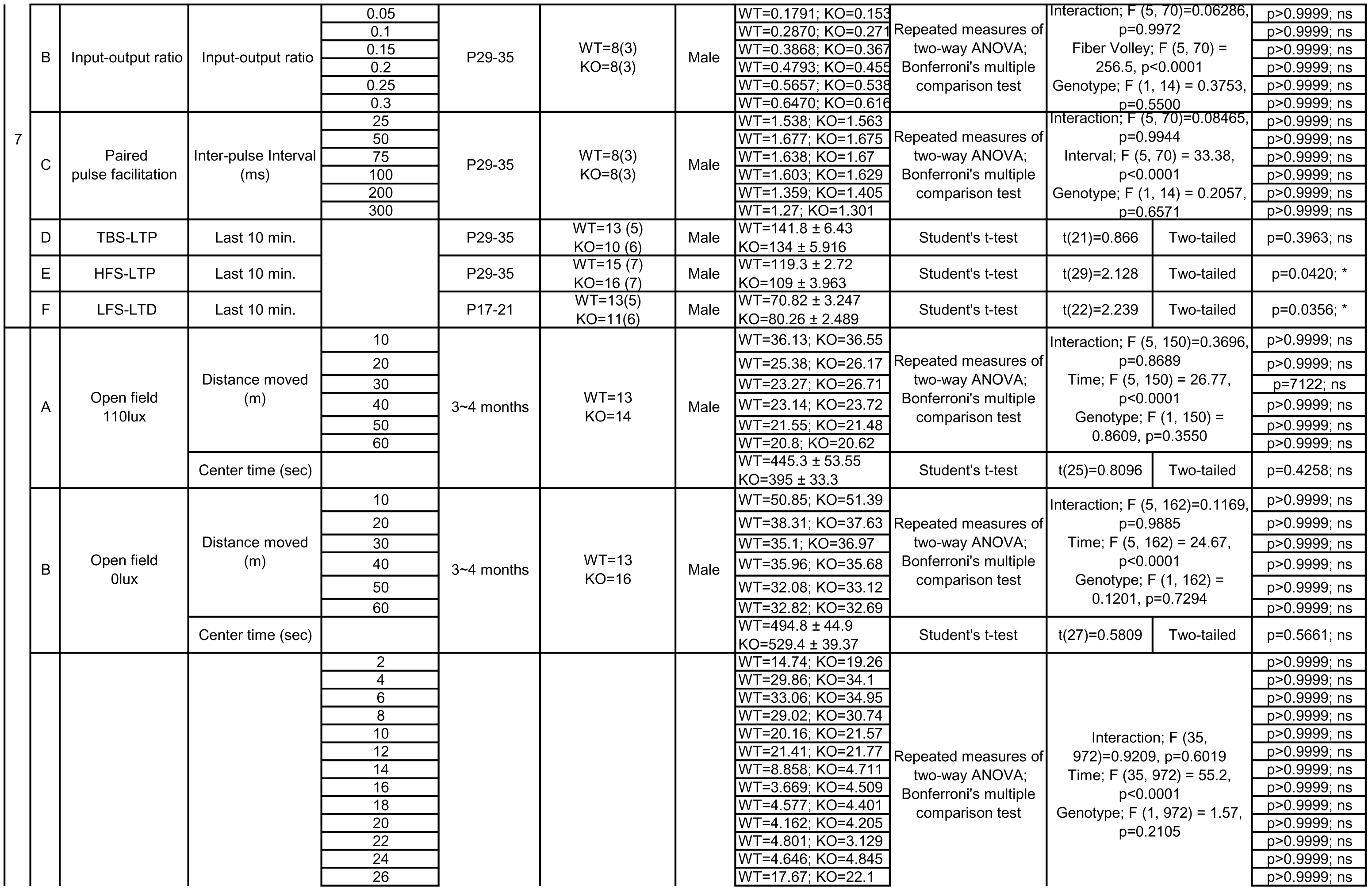
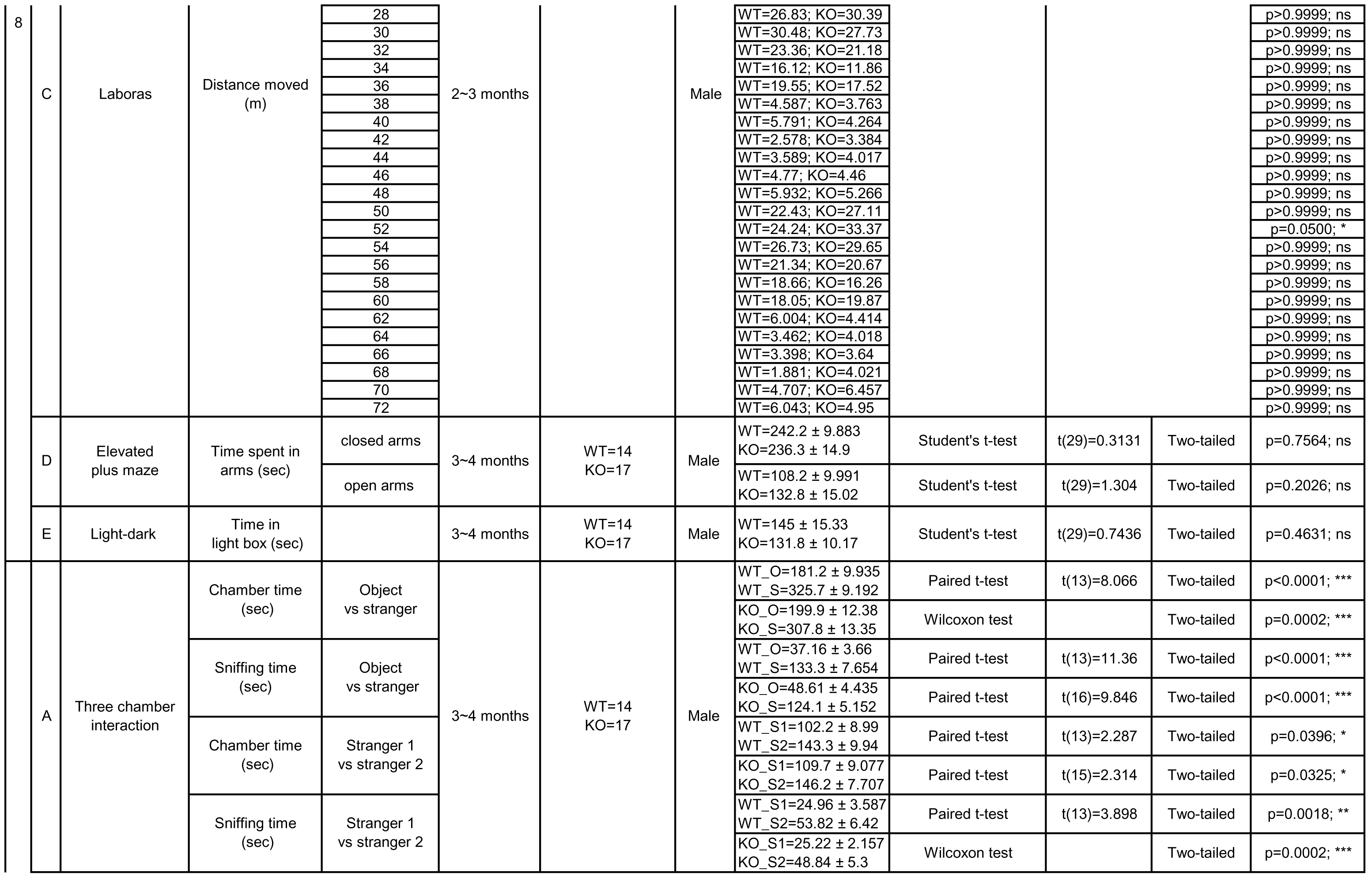
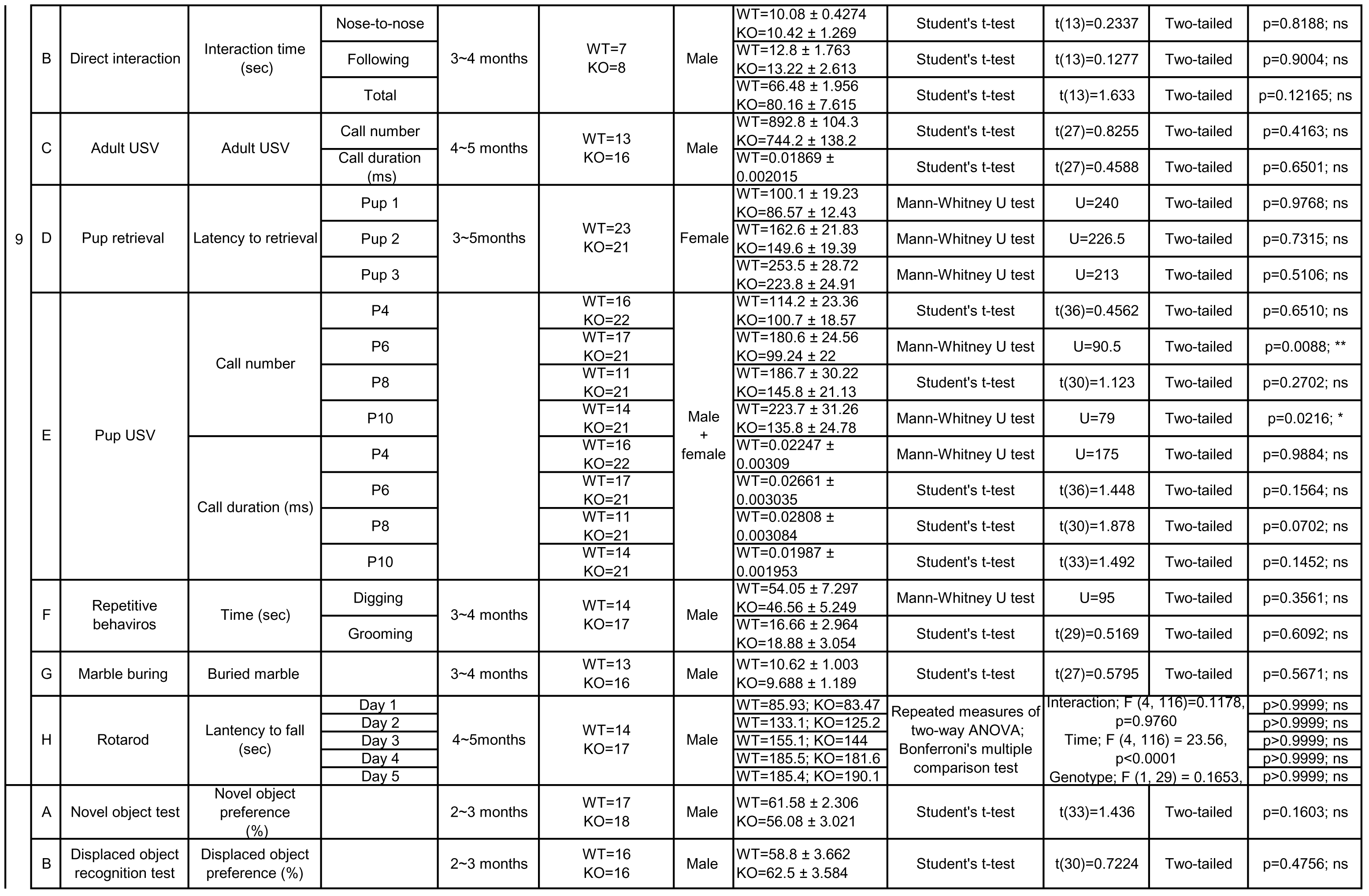
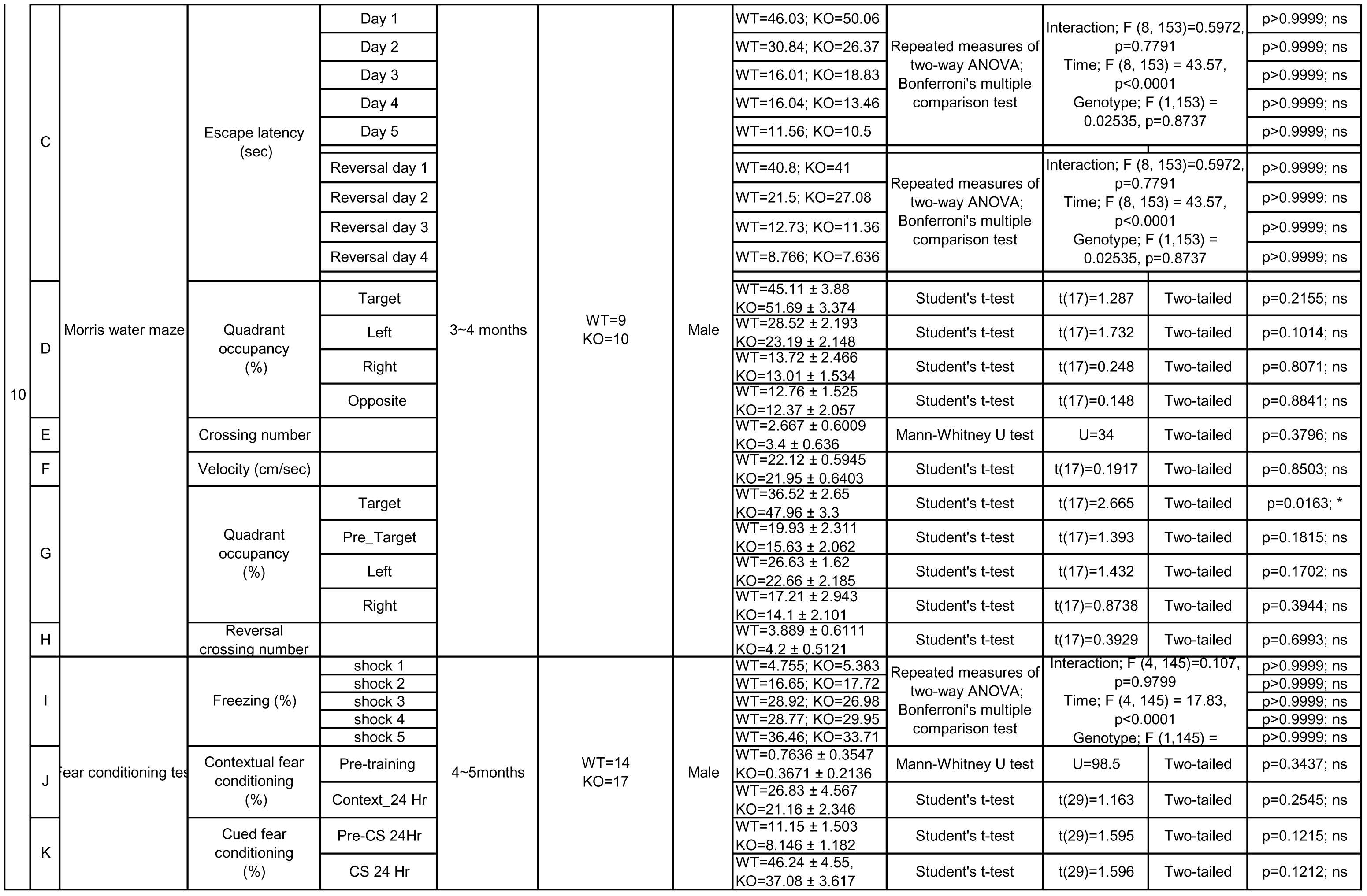
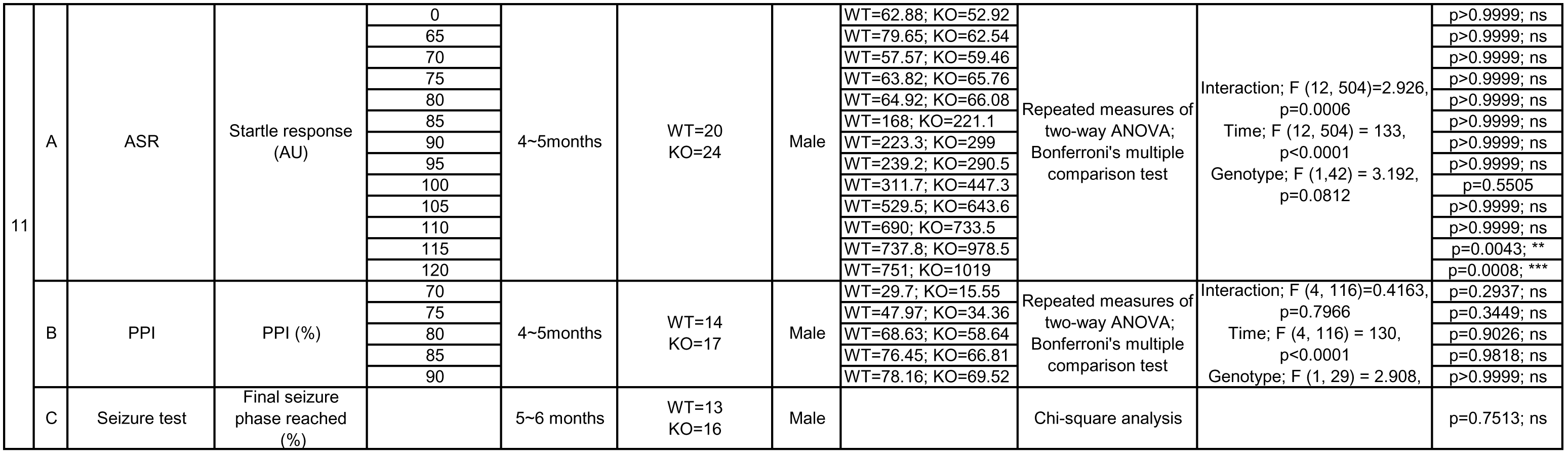
Details of animals used and statistical results.

## Results

### Generation and characterization of *Lrfn2*^*–/–*^ mice

To generate *Lrfn2^-/-^* mice, we used a mouse embryonic stem cell line that lacks exon 2 of the *Lrfn2* gene encoding most of the extracellular region of SALM1 (**figure1A**). This deletion was confirmed by polymerase chain reaction (PCR) genotyping and quantitative reverse transcription (RT)-PCR (**Figure 1B,C**). Immunoblot analyses using an anti-SALM1 antibody raised against the last 30 amino acid residues of the protein confirmed the lack of SALM1 protein in the *Lrfn2^-/-^* hippocampus (**Figure 1D**). These mice were born in normal Mendelian ratios, and did not exhibit any gross anatomical abnormalities in the brain (**Figure 1E**).

**Figure 1.**
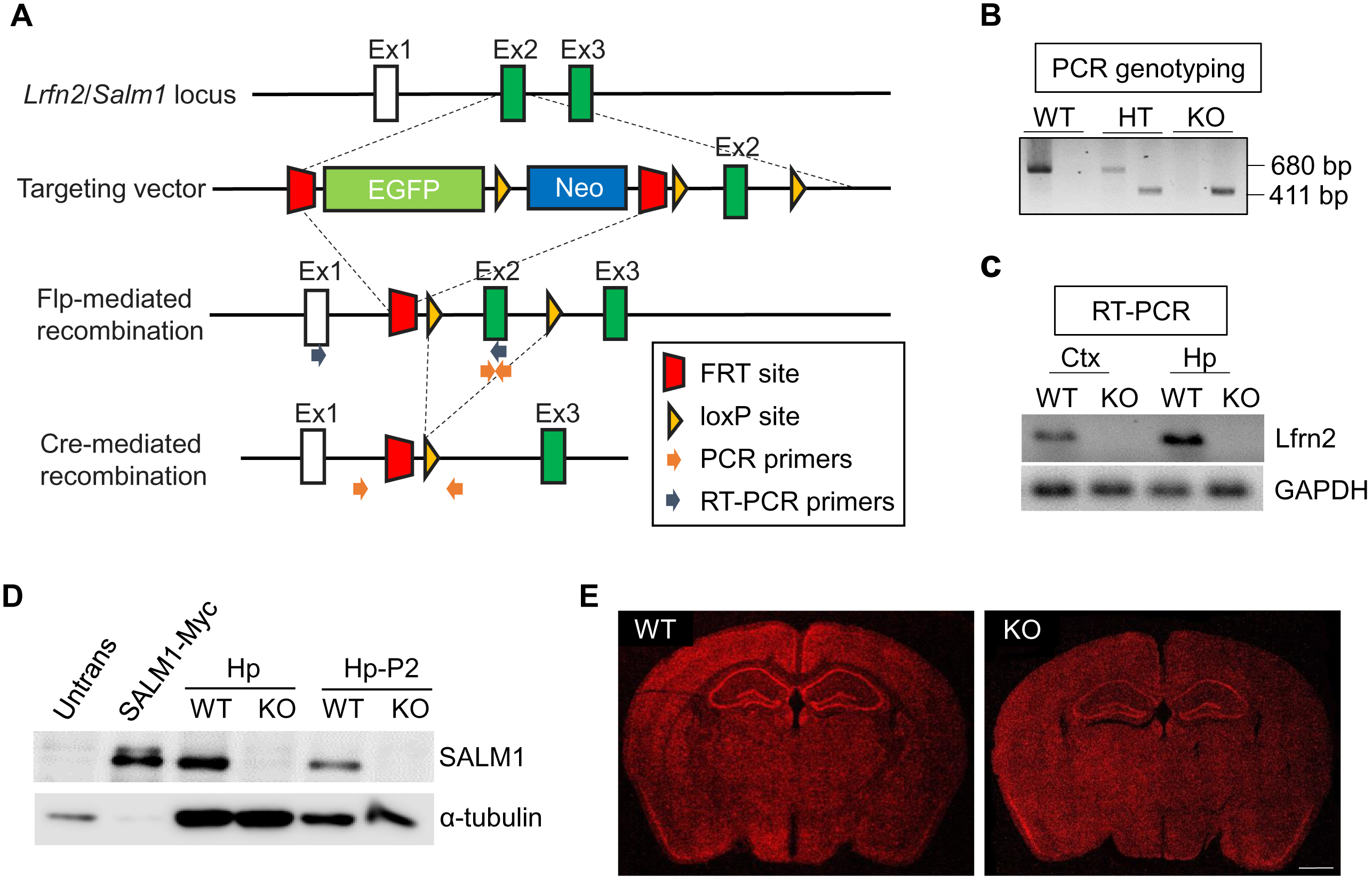
Generation and characterization of *Lrfn2*^*-/-*^ mice. (A) Strategy for the generation of *Lrfn2*^*-/-*^ mice. (B) PCR genotyping of *Lrfn2*^*-/-*^ mice. WT, wild-type; HT, heterozygous; KO, homozygous knockout. (C) Confirmation of *Lrfn2* exon 2 deletion by qRT-PCR analysis of whole-brain mRNAs (P31). (D) Lack of LRFN2 protein expression in the *Lrfn2*^*-/-*^ brain (P21), determined by immunoblot analysis of hippocampal lysates (total [Hp] and crude synaptosomal [Hp-P2]). Untrans, untransfected HEK293 cell lysates; SALM1-Myc, lysates of HEK293 cells transfected with SALM1-Myc. (E) Normal gross morphology of the *Lrfn2*^*-/-*^ brain (P28), as shown by staining for NeuN (a neuronal marker). Scale bar, 1 mm.

### Distribution patterns of SALM1 mRNA and protein

We next determined the distribution pattern of SALM1 mRNA at various developmental stages (embryonic day 18, postnatal day [P] 0, P7, P14, P21, and P42) by in situ hybridization in mouse brain slices. SALM1 mRNA signals in sagittal and horizontal sections, revealed by two independent probes, were relatively strong in cortical areas of the brain until P0, and gradually increased in other brain regions across postnatal developmental stages (**Figure 2**).

**Figure 2.**
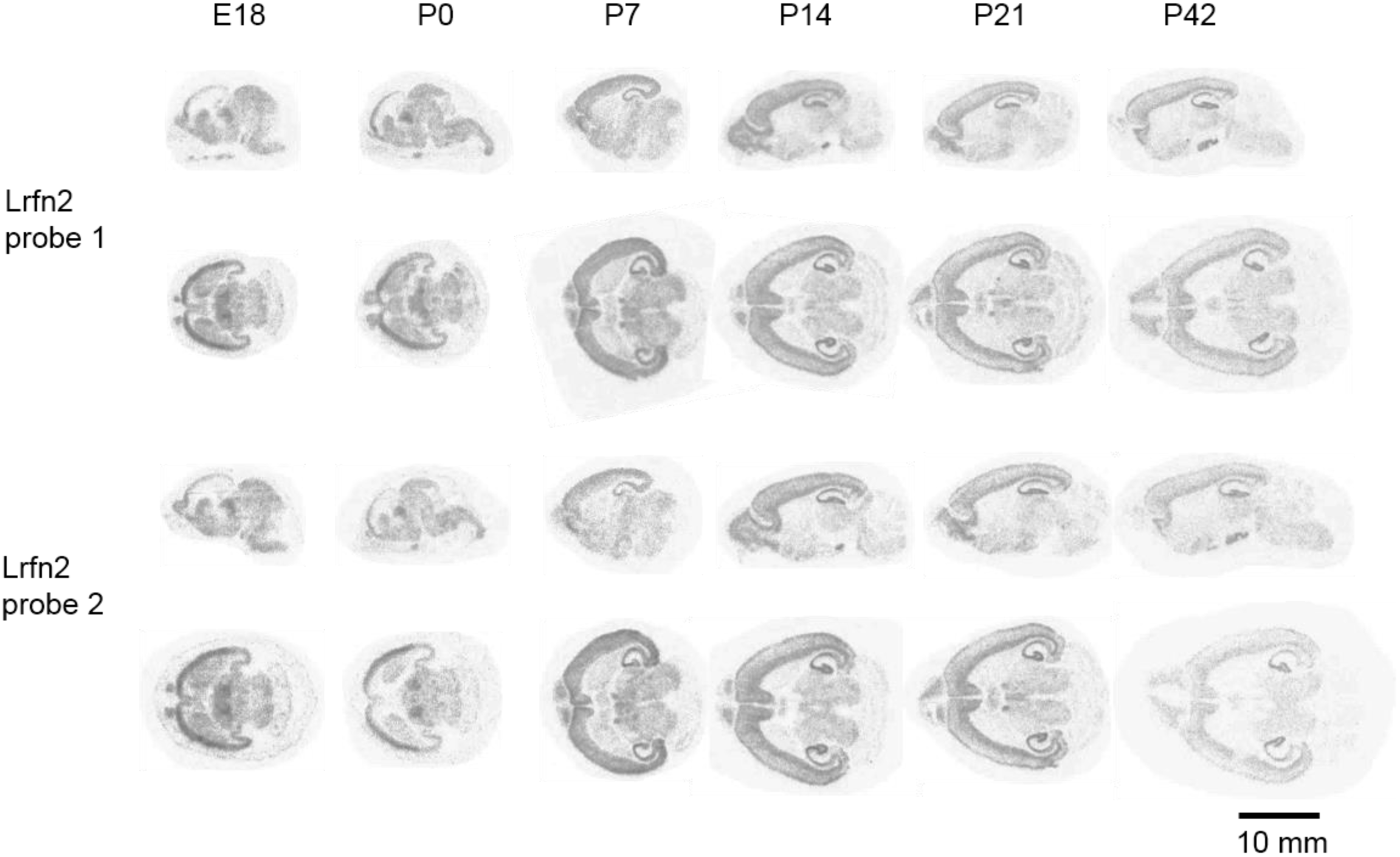
SALM1 mRNA distribution pattern. (A and B) In situ hybridization analysis of SALM1/Lrfn2 mRNA expression in mouse brain slices at different developmental stages using two independent probes against two different regions of SALM1/Lrfn2 mRNA. E, embryonic day; P, postnatal day. Scale bar, 10 mm.

To determine the distribution pattern of SALM1 protein in the brain, we used another line of transgenic mice in which the entire open reading frame of the *Lrfn2* gene was replaced with a LacZ cassette (KOMP VG15208; termed *Lrfn2-LacZ* mice) (Valenzuela et al., 2003). X-gal staining of coronal and sagittal sections from *Lrfn2-LacZ* mice (P46; male HT) revealed that SALM1 protein is highly expressed in various brain regions, including the cortex, hippocampus, amygdala, thalamus, and hypothalamus (**Figure 3A,B**). Notably, SALM1 protein was more abundant in layers II/III and VI of the cortex relative to the middle layers, and in CA1 and CA3 regions of the hippocampus relative to the dentate gyrus. In contrast, SALM1 protein was minimally detected in the striatum and cerebellum.

**Figure 3.**
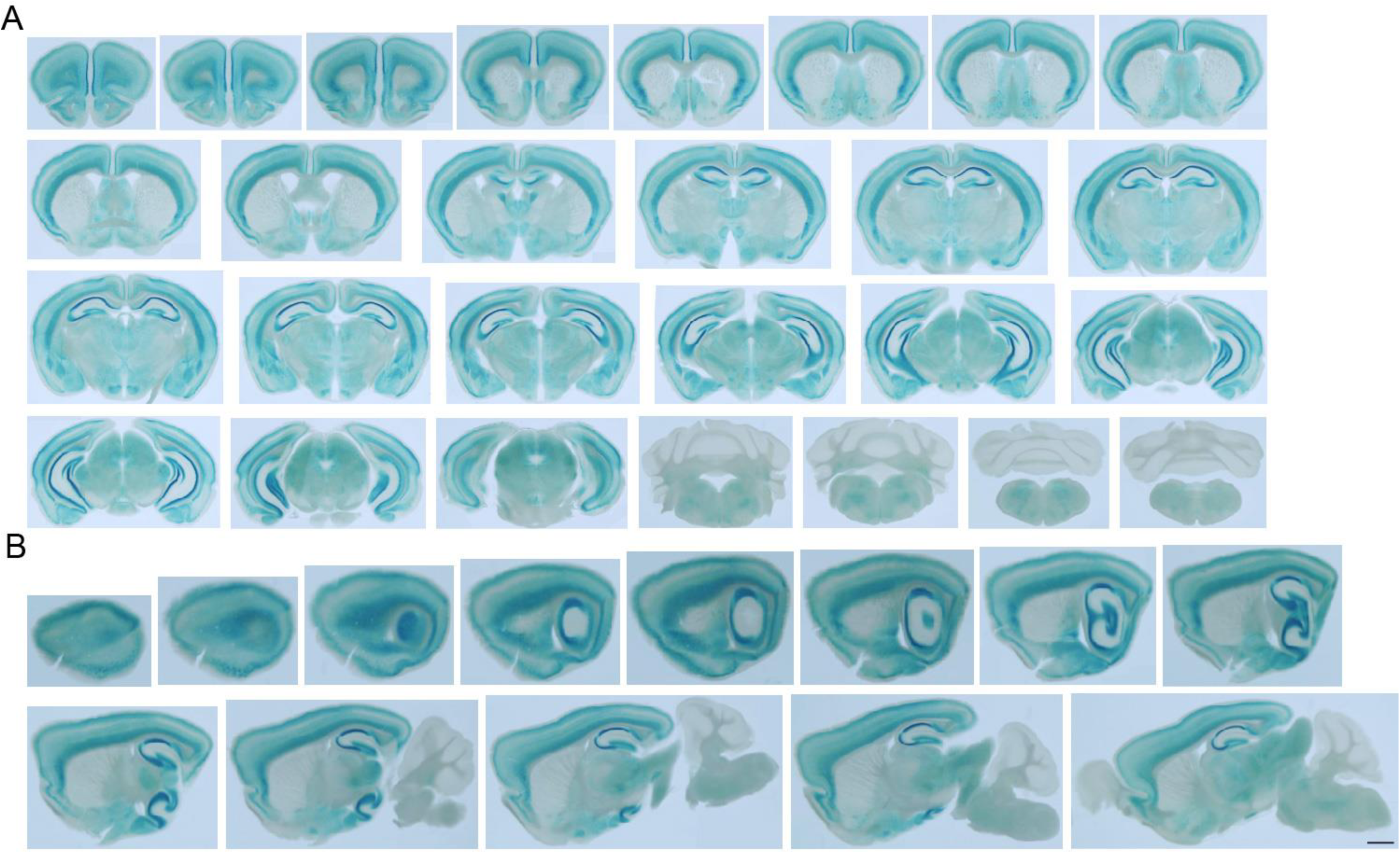
SALM1 protein distribution pattern. (A and B) Distribution pattern of SALM1 protein, determined by X-gal staining of coronal and sagittal sections of brain from male heterozygous (*Lrfn2*^*+/–*^) mice (P46). Scale bar, 1 mm.

### Suppressed excitatory and inhibitory synaptic transmission in *Lrfn2*^*-/-*^ mice

To determine whether *Lrfn2* deletion has any effects on synapse development and function, we first determined spontaneous excitatory and inhibitory synaptic transmission in the CA1 region of the hippocampus, where SALM1 is highly expressed. We found that both the frequency and amplitude of miniature excitatory postsynaptic currents (mEPSCs) in *Lrfn2*^*-/-*^ CA1 pyramidal neurons (P21–23) were comparable to those from wild-type (WT) mice (**Figure 4A**).

**Figure 4.**
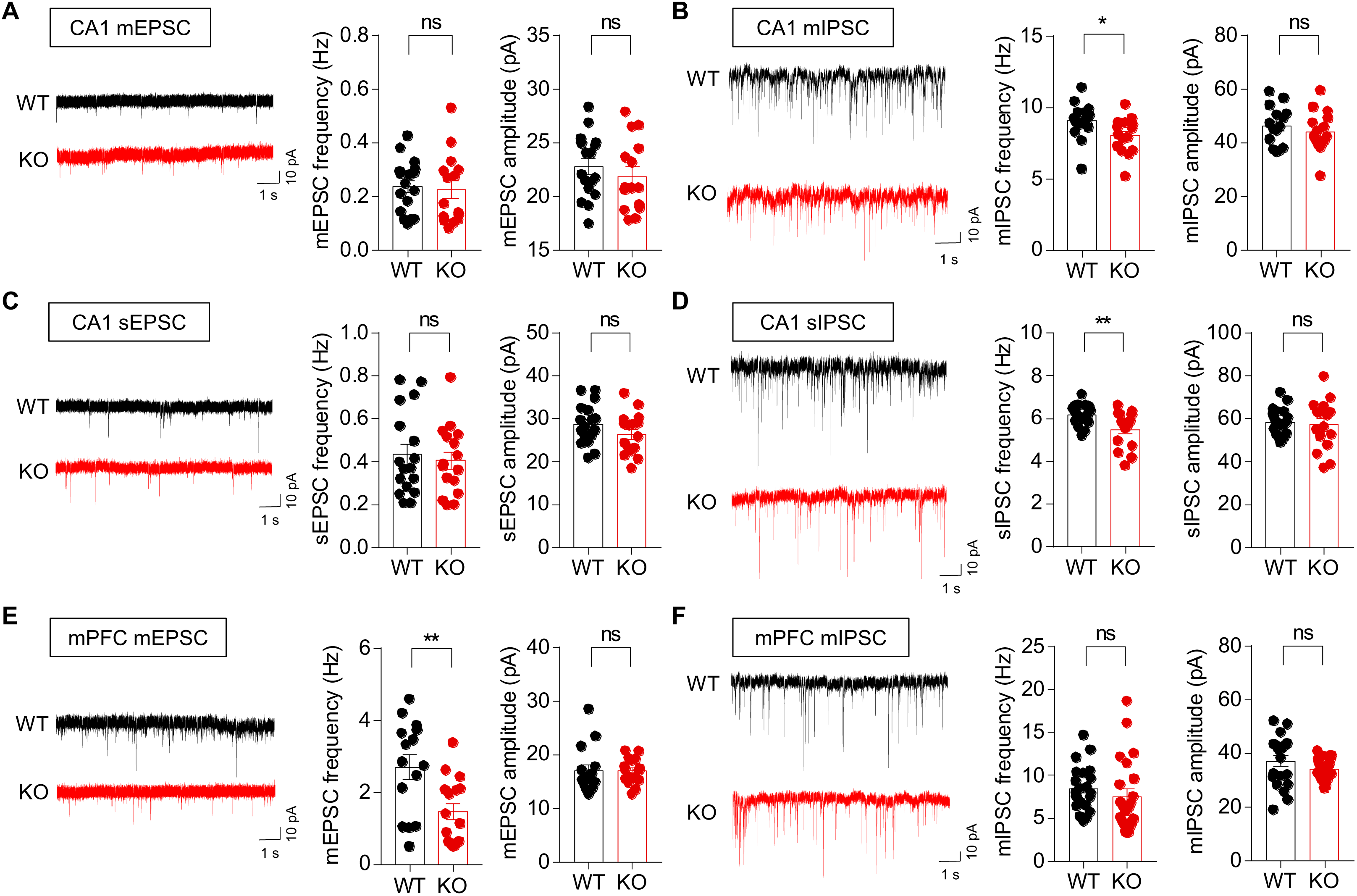
Suppressed excitatory and inhibitory synaptic transmission in *Lrfn2*^*-/-*^ mice. (A) mEPSCs measured in WT and *Lrfn2*^*-/-*^ CA1 pyramidal neurons (P20–23). n = 17 cells from 4 WT mice and 15 cells from 6 KO mice, ns, not significant, Student’s ttest. (B) mIPSCs in WT and *Lrfn2*^*-/-*^ CA1 pyramidal neurons (P20–23). n = 15, 3 for WT and 15, 3 for KO., *p < 0.05, ns, not significant, Mann-Whitney U test and Student’s t-test. (C) sEPSCs in WT and *Lrfn2*^*-/-*^ CA1 pyramidal neurons (P20–23). n = 17, 5 for WT and 16, 5 for KO, ns, not significant, Student’s t-test. (D) sIPSCs in WT and *Lrfn2*^*-/-*^ CA1 pyramidal neurons (P20–23). n = 18, 3 for WT and 16, 3 for KO, **p < 0.01, ns, not significant, Student’s t-test. (E) mEPSCs in WT and *Lrfn2*^*-/-*^ mPFC prelimbic layer II/III pyramidal neurons (P20–23). n = 15, 3 for WT and 16, 3 for KO, **p < 0.01, ns, not significant, Mann-Whitney U test and Student’s t-test. (F) mIPSCs in WT and *Lrfn2*^*-/-*^ mPFC prelimbic layer II/III pyramidal neurons (P20–23). n = 17, 3 for WT and 19, 3 for KO, **p < 0.01, ns, not significant, Mann-Whitney U test and Student’s t-test.

In contrast, the frequency, but not amplitude, of miniature inhibitory postsynaptic currents (mIPSCs) was significantly reduced in *Lrfn2^-/-^* CA1 pyramidal neurons (P20–23) (**Figure 4B**). Similar results were obtained for spontaneous EPSCs (sEPSCs) and sIPSCs, revealing a specific decrease in sIPSC frequency (**Figure 4C,D**); these latter recordings were obtained in the absence of tetrodotoxin to allow action potential firing and network activities. These results suggest that a SALM1 deficiency leads to a decrease in the frequency of inhibitory, but not excitatory, synaptic transmission in the hippocampal CA1 region, and that this decrease is not compensated by network activities.

To further test if *Lrfn2* deletion leads to similar changes in other brain regions, we measured mEPSCs and mIPSCs from layer II/III pyramidal neurons in the prelimbic region of the medial prefrontal cortex (mPFC). We found that the frequency, but not amplitude, of mEPSCs, was significantly reduced, whereas mIPSCs were unaltered (**Figure 4E,F**), a finding that contrasts with the results obtained in the hippocampus. These results collectively suggest that *Lrfn2* deletion suppresses the frequency of excitatory synaptic transmission in the mPFC, but the frequency of inhibitory transmission in the hippocampus.

### Decreased inhibitory synapse density in the *Lrfn2*^*-/-*^ hippocampus

To further understand the mechanism underlying the suppressed synaptic transmission in *Lrfn2*^*-/-*^ mice, we analyzed the density and morphology of excitatory and inhibitory synapses using electron microscopy (EM). Morphologically, excitatory and inhibitory synapses were defined by postsynaptic densities (PSDs) apposed to presynaptic axon terminals and PSDs apposed to GABA immuno-positive inhibitory axon terminals, respectively. We found no significant changes in the density of excitatory synapses in the CA1 stratum radiatum region of the *Lrfn2*^*-/-*^ hippocampus compared with WT controls (**Figure 5A,B**). In addition, there were no changes in the length, thickness, or perforation of *Lrfn2*^*-/-*^ PSDs.

**Figure 5.**
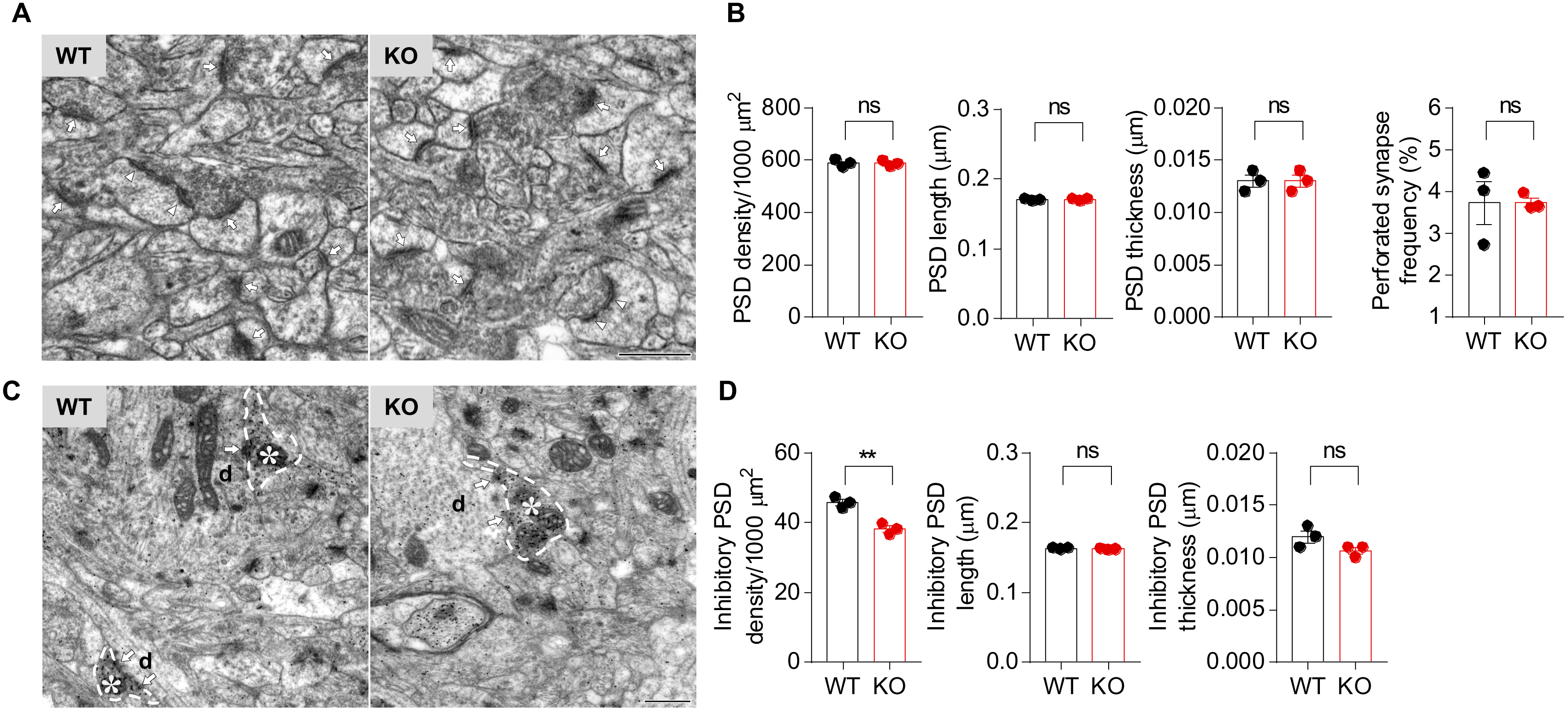
Decreased inhibitory synapse density in the *Lrfn2*^*-/-*^ hippocampus. (A and B) Decreased density of inhibitory, but not excitatory, synapses in the stratum radiatum region of the CA1 region in *Lrfn2*^*-/-*^ mice (P21), as determined by EM analysis. Excitatory synapses were defined by PSD structures apposed to axon terminals (arrows and arrowheads for non-perforated and perforated PSD, respectively), and inhibitory synapses were defined by PSDs apposed to presynaptic axon terminals with immunopositive GABA signals (arrows). Asterisks, GABA immune-positive axon terminals; d, postsynaptic dendrites. Scale bar, 500 nm. n = 3 mice for WT and KO, **p < 0.01, ns, not significant, Student’s t-test.

In contrast, an analysis of inhibitory synapses indicated a significant decrease in the density, but not the length or thickness, of PSDs (**Figure 5C,D**). These results suggest that a SALM1 deficiency leads to a decrease in the density of inhibitory, but not excitatory, synapses in the hippocampus. Taken together with the synaptic transmission results, these findings suggest that the reduced inhibitory synapse number contributes to the reduced frequency of inhibitory synaptic transmission.

### SALM1 expression in both glutamatergic and GABAergic neurons

The decrease in inhibitory synapse density associated with a SALM1 deficiency might be attributable to cell-autonomous changes in CA1 pyramidal neurons and/or alterations in presynaptic GABAergic neurons that express SALM1 protein. To address this question, we performed double-immunofluorescence in situ hybridization experiments for SALM1/Lrfn2 and the glutamatergic neuron marker, Vglut1/2 (vesicular glutamate transporter-1/2) and GABAergic neuron marker Gad1/2 (glutamate decarboxylase-1/2). Signals for SALM1/Lrfn2 mRNA were detected strongly in both hippocampal and cortical areas (**Figure 6**), where they were detected in cell bodies positive for Vglut1/2 as well as Gad1/2 (**Figure 6**). These results suggest that SALM1/Lrfn2 is expressed in both glutamatergic and GABAergic neurons in the cortex and hippocampus.

**Figure 6.**
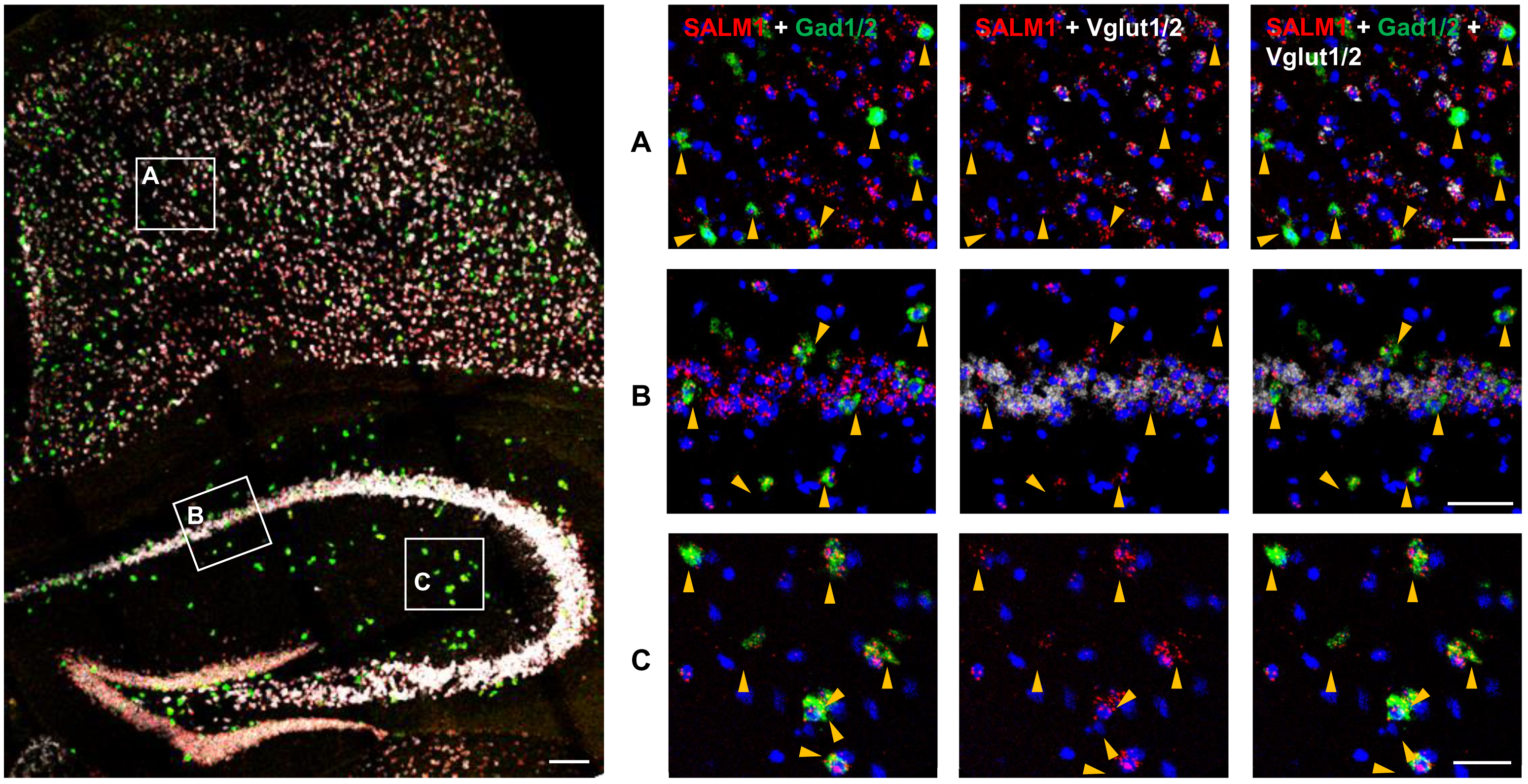
*Lrfn2* mRNA is detected in both glutamatergic and GABAergic neurons. Expression of SALM1/Lrfn2 mRNA in glutamatergic and GABAergic neurons, was determined by fluorescence in situ hybridization. Coronal sections from male mouse brains (P21, male, 8 weeks) were triply stained for SALM1/Lrfn2, Vglut1/2 (glutamatergic neuron markers) and Gad1/2 (GABAergic neuron markers), and counterstained with the nuclear dye DAPI. A mixture of two probes (Vglut1 + Vglut2, or Gad1 + Gad2) was used to label all glutamatergic or GABAergic neurons. The indicated cortical and hippocampal regions in the image at left were enlarged (right panels) to highlight the expression of Lrfn2 mRNA in both glutamatergic and GABAergic neurons; Lrfn2 expression in GABAergic neurons were further highlighted by arrows. Scale bar, 50 μm.

### Increased NMDA/AMPA ratio and suppressed NMDAR-dependent synaptic plasticity in the *Lrfn2*^*-/-*^ hippocampus

Although we found that a SALM1 deficiency has no effect on excitatory synapse density or spontaneous synaptic transmission in the hippocampus, given that SALM1 forms a complex with PSD-95 and NMDARs in vitro and in vivo (Ko et al., 2006; Morimura et al., 2006; Wang et al., 2006), it is possible that a SALM1 deficiency might alter other aspects of excitatory synapse development and function. To test this possibility, we first measured the ratio of evoked NMDAR‐ and AMPAR-mediated EPSCs (NMDA/AMPA ratio). Interestingly, these experiments revealed an increase in the NMDA/AMPA ratio at *Lrfn2*^*-/-*^ Schaffer collateral-CA1 pyramidal (SC-CA1) synapses (**Figure 7A**). In contrast, there were no changes in basal excitatory synaptic transmission (input-output relationship) or paired-pulse facilitation (**Figure 7B,C**). These results suggest that it is unlikely that AMPAR-mediated synaptic transmission or presynaptic release probability are changed in *Lrfn2*^*-/-*^ SC-CA1 synapses, and that the increased NMDA/AMPA ratio in these synapses may be attributable to an increase in NMDAR-mediated EPSCs.

**Figure 7.**
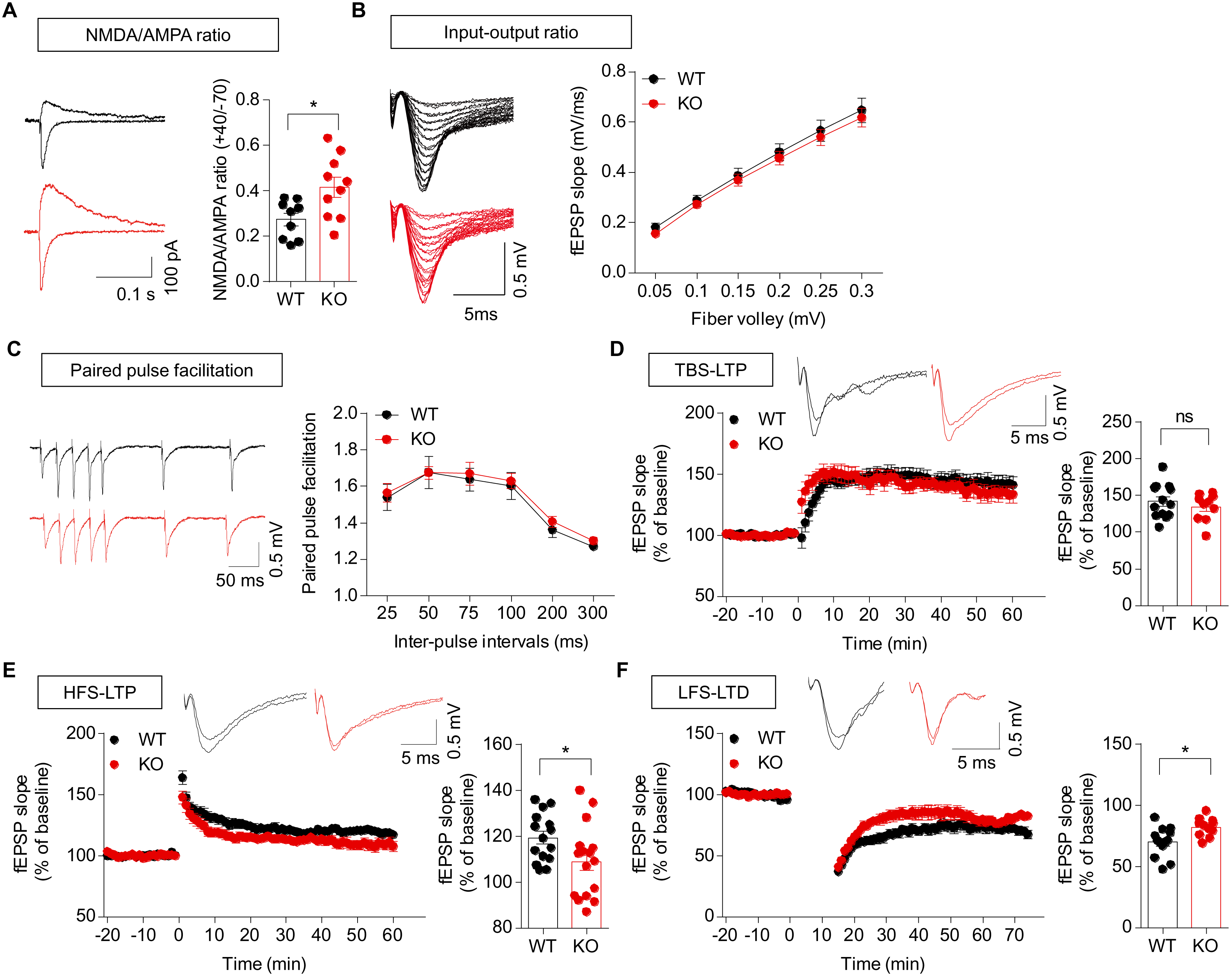
Increased NMDA/AMPA ratio and suppressed NMDAR-dependent synaptic plasticity in the *Lrfn2*^*-/-*^ hippocampus. (A) Increased ratio of NMDAR/AMPAR-mediated synaptic transmission at *Lrfn2*^*-/-*^ SC-CA1 synapses (P21–24), measured as AMPA and NMDA EPSCs evoked at holding potentials of –70 and +40 mV, respectively. n = 9 slices from 8 mice for WT and 10 slices from 8 mice for KO, *p < 0.05, Student’s t-test. (B) Normal basal transmission at *Lrfn2*^*-/-*^ SC-CA1 synapses (P30–35), as shown by the input-output relationship between fiber volley and fEPSP slopes. n = 8, 3 for WT and KO, repeated measures two-way ANOVA. (C) Normal paired pulse facilitation at *Lrfn2*^*-/-*^ SC-CA1 synapses (P30–35), as shown by the relationship between inter-pulse intervals and fEPSP slopes. n = 8, 3 for WT and KO, repeated measures two-way ANOVA. (D) Normal TBS-LTP at *Lrfn2*^*-/-*^ SC-CA1 synapses (P28–36). Bar graphs represent average values during the last 10 minutes. n = 13, 5 for WT and 10, 6 for KO, ns, not significant, Student’s t-test. (E) Suppressed HFS-LTP at *Lrfn2*^*-/-*^ SC-CA1 synapses (P28–36). Bar graphs represent average values during the last 10 minutes. n = 15, 7 for WT and 16, 7 for KO, *p < 0.05, Student’s t-test. (F) Suppressed LFS-LTD at *Lrfn2*^*-/-*^ SC-CA1 synapses (P16–20). Bar graphs represent average values during the last 10 minutes. n = 13, 5 for WT and 11, 6 for KO, *p < 0.05, Student’s t-test.

Given that NMDAR function regulates synaptic plasticity, affecting both long-term potentiation (LTP) and long-term depression (LTD), we measured these parameters at *Lrfn2*^*-/-*^ SC-CA1 synapses. Contrary to our initial expectation that NMDAR-dependent synaptic plasticity would be increased, we found that LTP induced by theta burst stimulation (TBS-LTP) at *Lrfn2*^*-/-*^ synapses was normal (**Figure 7D**). In contrast, LTP induced by high-frequency stimulation (HFS-LTP) and LTD induced by low-frequency stimulation (LFS-LTD) were significantly decreased at *Lrfn2*^*-/-*^ synapses (**Figure 7E,F**). These results collectively suggest that a SALM1 deficiency suppresses specific forms of NMDAR-dependent LTP and LTD without affecting basal excitatory transmission or presynaptic neurotransmitter release.

### Normal locomotion and anxiety-like behavior in *Lrfn2*^*-/-*^ mice

To determine the impacts of SALM1 deletion on behaviors, we subjected *Lrfn2*^*-/-*^ mice to a battery of behavioral tests. *Lrfn2*^*-/-*^ mice showed normal locomotor activity in the open field test under both bright-light (110 lux) and light-off (0 lux) conditions (**Figure 8A,B**). Continuous monitoring of mouse movements for 3 days in a home cage-like environment (Laboras cage) revealed no abnormalities in the locomotor activity of *Lrfn2*^*-/-*^ mice (**Figure 8C**).

**Figure 8.**
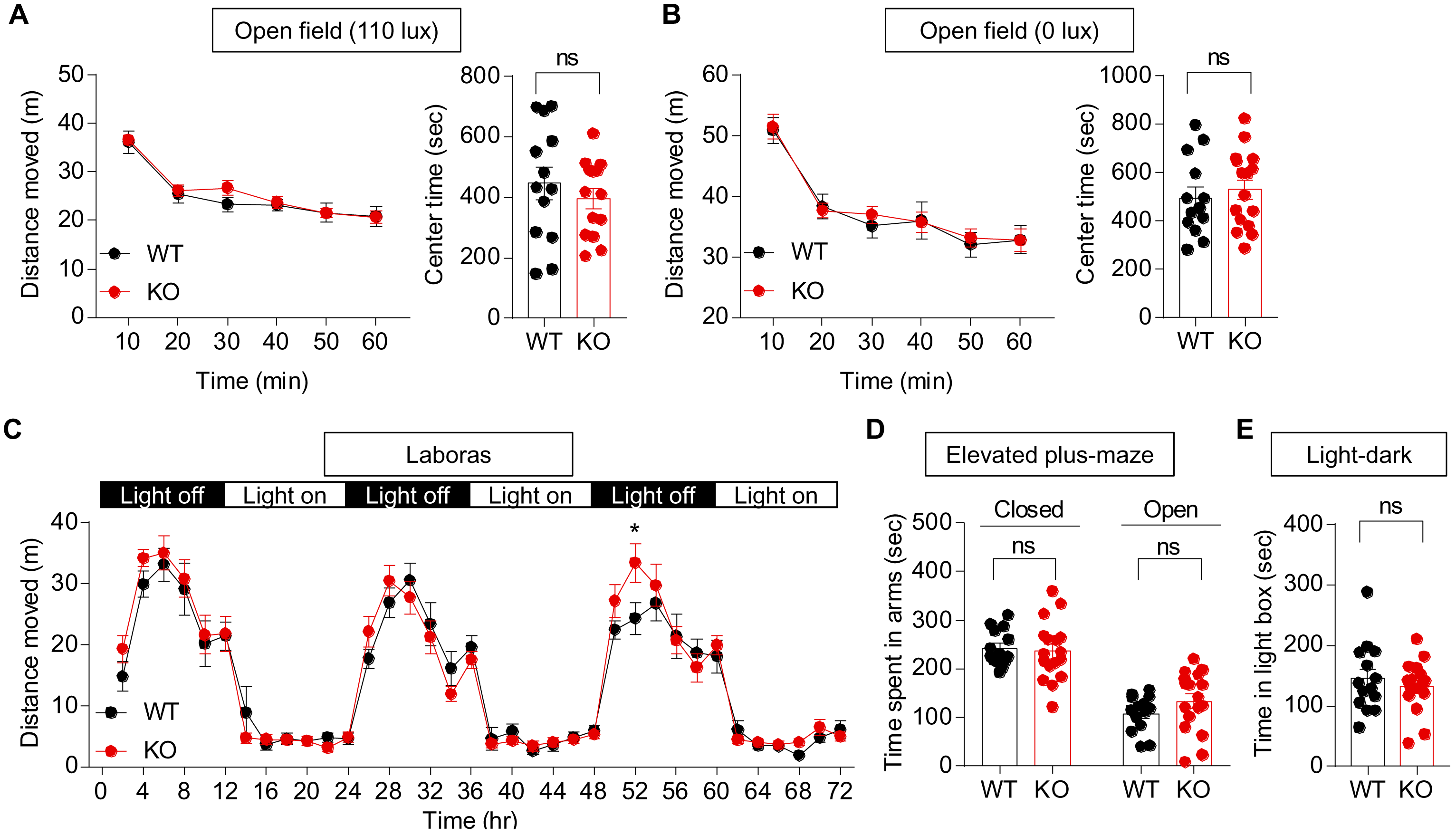
Normal locomotion and anxiety-like behavior in *Lrfn2*^*-/-*^ mice. (A and B) Normal locomotor activity and anxiety-like behavior of *Lrfn2*^*-/-*^ mice (3 months) in open field tests at two different light intensities (110 and 0 lux), as shown by distance moved and time spent in the center region of the open field arena. n = 13 mice for WT and 14 mice for KO (110 lux), and 13 for WT and 16 for KO (0 lux), ns, not significant, repeated measures two-way ANOVA and Student’s t-test. (C) Normal locomotor activity of *Lrfn2*^*-/-*^ mice (2 months) in Laboras cages, where mouse movements were monitored for three consecutive days. n = 13 for WT and 16 for KO, ns, not significant, repeated measures two-way ANOVA. (D) Normal anxiety-like behavior of *Lrfn2*^*-/-*^ mice (4 months) in the elevated plus-maze test. n = 14 for WT and 17 for KO, ns, not significant, Student’s t-test. (E) Normal anxiety-like behavior of *Lrfn2*^*-/-*^ mice (4 months) in the light-dark test. n = 14 for WT and 16 for KO, ns, not significant, Student’s t-test.

*Lrfn2*^*-/-*^ mice showed normal anxiety-like behaviors in the elevated plus-maze (**Figure 8D**), the light-dark test (**Figure 8E**), and the open field test (center time) (**Figure 8A,B**). These results collectively suggest that a SALM1 deficiency has minimal effects on locomotor activity and anxiety-like behavior.

### Suppressed USVs in *Lrfn2*^*-/-*^ pups, but normal social interaction and repetitive behaviors in adult *Lrfn2*^*-/-*^ mice

We next examined social behaviors. *Lrfn2*^*-/-*^ mice displayed normal social interaction and social novelty recognition in the three-chamber test (**Figure 9A**) and the direct social-interaction test (**Figure 9B**). Measurements of USVs revealed that female encounters elicited normal levels of USVs in adult male *Lrfn2*^*-/-*^ mice (**Figure 9C**). In addition, adult female *Lrfn2*^*-/-*^ mice retrieved pups to an extent similar to that of WT females (**Figure 9D**). In contrast, USVs emitted by *Lrfn2^-/-^* pups separated from their mothers were diminished, as shown by the number of calls and individual call duration (**Figure 9E**).

**Figure 9.**
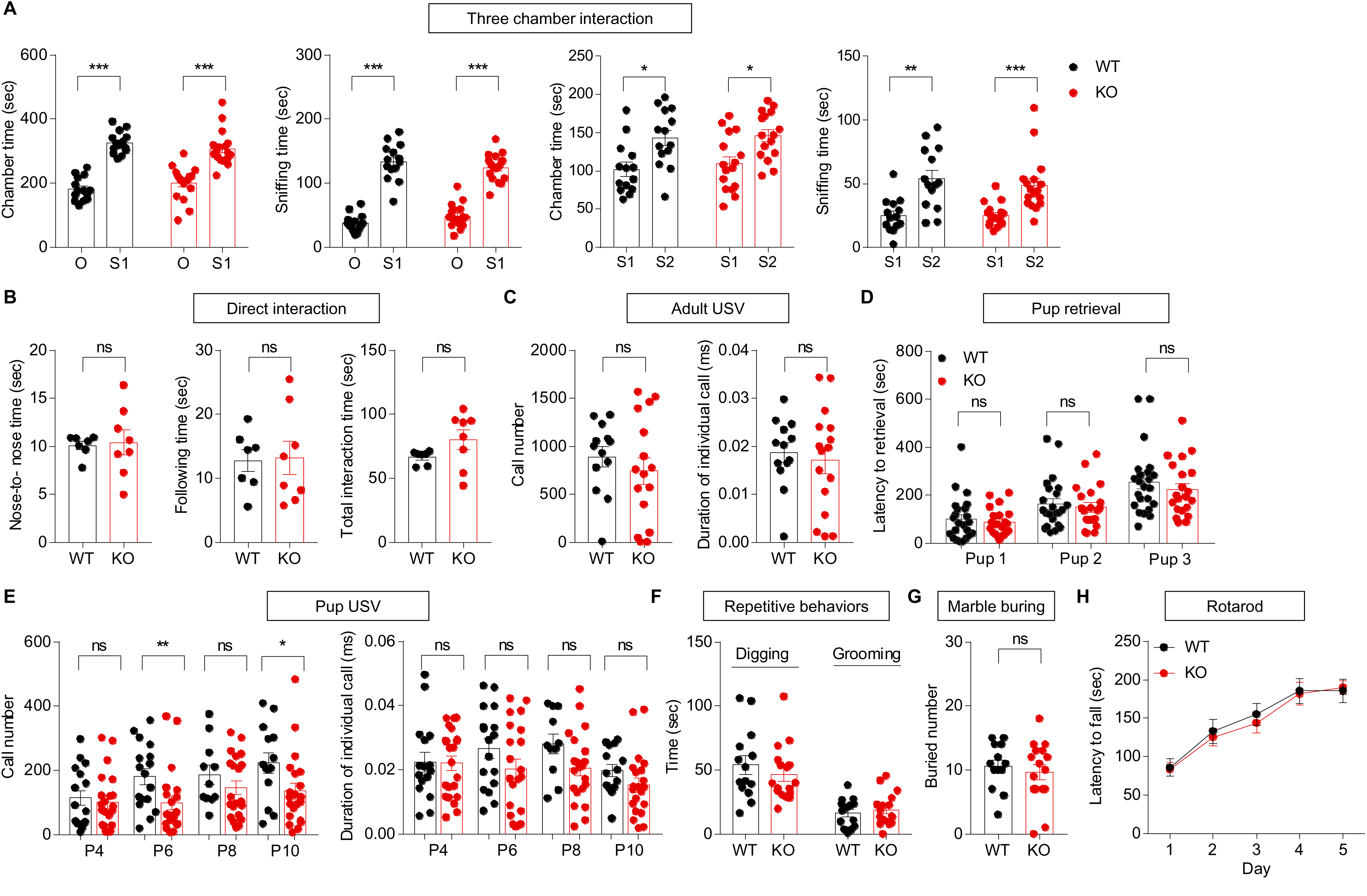
Suppressed USVs in pups, but normal social interaction, communication, and repetitive behaviors in adult *Lrfn2*^*-/-*^ mice. (A) Normal social approach and social novelty recognition in *Lrfn2*^*-/-*^ mice (3–4 months) in the three-chamber test, as shown by time spent in the chamber and sniffing. n = 14 mice for WT and 17 mice for KO, *p < 0.05, **p < 0.01, ***p < 0.001, Paired t-test and Wilcoxon test. (B) Normal social interaction in *Lrfn2*^*-/-*^ mice (3–4 months) in the direct social-interaction test. n = 7 WT mouse pairs and 8 KO mouse pairs, ns, not significant, Student’s t-test. (C) Normal USVs of male *Lrfn2*^*-/-*^ mice (4–5 months) induced by encounter with a female. n = 13 mice for WT and 16 mice for KO, ns, not significant, Student’s t-test. (D) Normal pup retrieval in female *Lrfn2*^*-/-*^ mice (3–5 months) induced by WT pups (P1) separated from their mothers, as shown by the time taken to retrieve first, second, and third pups. n = 23 mice for WT and 21 mice for KO, ns, not significant, Mann-Whitney U test. (E) Suppressed USVs in *Lrfn2*^*-/-*^ pups (P4, P6, P8, and P10) separated from their mothers, as shown by number of calls and individual call duration. For P4, n = 16 mice for WT and 22 mice for KO; P6, n = 17 for WT and 21 for KO; P8, n = 11 for WT and 21 for KO; P10, n = 14 for WT and 21 for KO, *p < 0.05, ns, not significant, Student’s t-test and Mann-Whitney U test. (F and G) Normal repetitive behaviors of *Lrfn2*^*-/-*^ mice (3–4 months), as shown by self-grooming, digging, and marble burying. For digging and grooming, n = 14 mice for WT and 17 mice for KO; marble burying, n = 13 for WT and 16 for KO, ns, not significant, Student’s t-test and Mann-Whitney U test. (H) Normal motor coordination of *Lrfn2*^*-/-*^ mice (4–5 months) in the rotarod test. n = 14 mice for WT and 17 mice for KO, repeated measures two-way ANOVA.

In tests measuring repetitive behaviors, *Lrfn2*^*-/-*^ mice showed normal levels of self-grooming, digging, and marble burying in home cages (**Figure 9F,G**). In addition, *Lrfn2*^*-/-*^ mice showed normal motor coordination and learning in the rotarod test (**Figure 9H**). Collectively, these results suggest that a SALM1 deficiency does not affect social interaction or repetitive behaviors, but does affect USVs, a form of social communication, in pups but not in adult mice.

### Normal learning and memory in *Lrfn2*^*-/-*^ mice

Turning to learning and memory behaviors, we found that *Lrfn2*^*-/-*^ mice performed normally in the novel object-recognition test and displaced object-recognition tests (**Figure 10A,B**). In the Morris water-maze test, *Lrfn2*^*-/-*^ mice showed normal levels of learning and memory in learning, probe, and reversal phases (**Figure 10C–H**). In fear-conditioning tests, in which mice were exposed to foot shocks in a spatial context combined with a tone, *Lrfn2*^*-/-*^ mice showed normal levels of freezing in the same spatial context 24 hours after acquisition of fear memory (**Figure 10I,J**). Freezing induced by the same sound cue 28 hours after fear memory acquisition was also normal in *Lrfn2*^*-/-*^ mice (**Figure 10K**). These results collectively suggest that a SALM1 deficiency has minimal effects on learning and memory behaviors.

**Figure 10.**
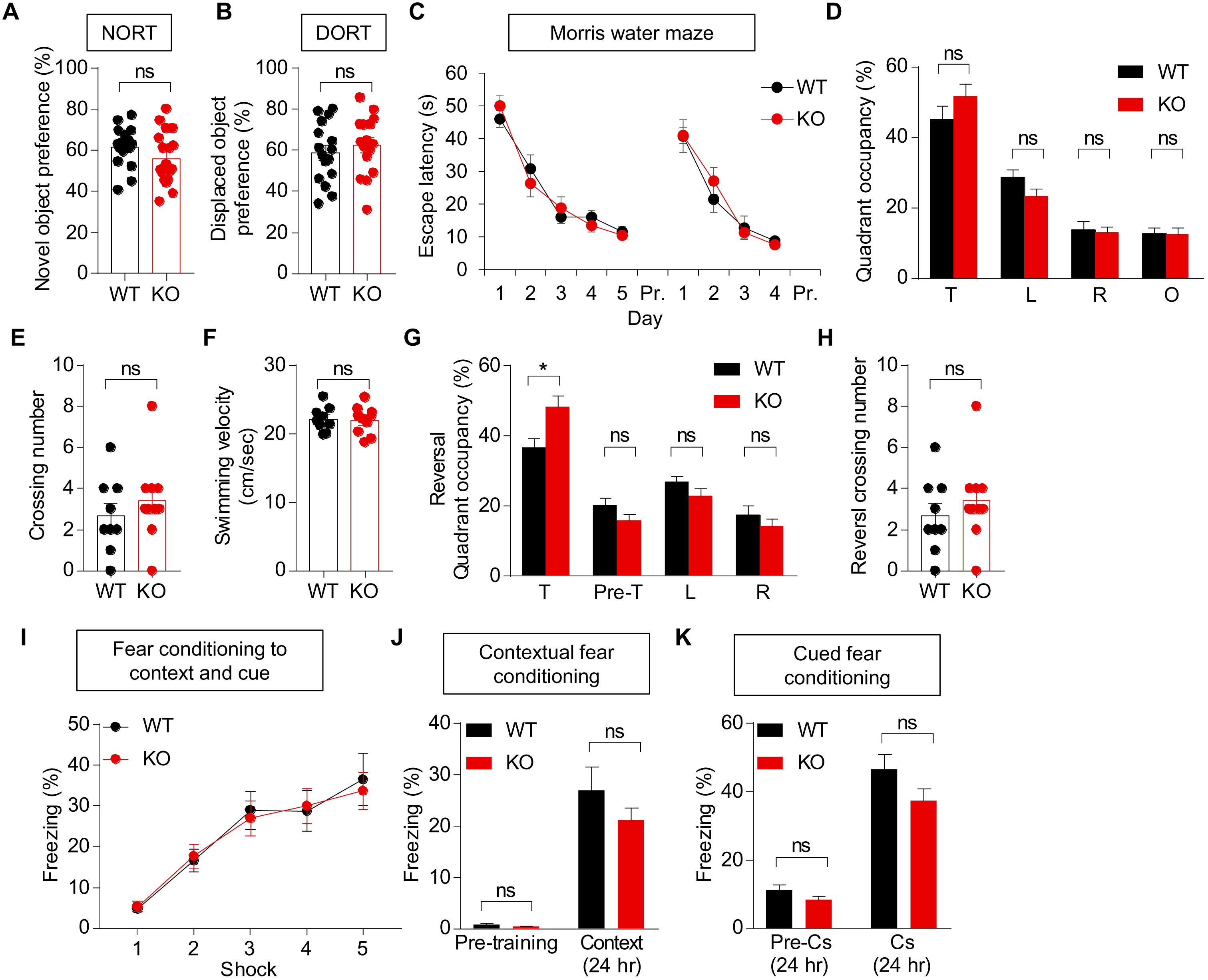
Normal learning and memory in *Lrfn2*^*-/-*^ mice. (A) Normal novel objection-recognition memory in *Lrfn2*^*-/-*^ mice (2–3 months). n = 17 mice for WT and 18 mice for KO, ns, not significant, Student’s t-test. (B) Normal displaced object-recognition memory in *Lrfn2*^*-/-*^ mice (2–3 months). n = 16 mice for WT and KO, ns, not significant, Student’s t-test. (C–H) Normal learning and memory in *Lrfn2*^*-/-*^ mice (3–4 months) in the learning, probe, and reversal phases of the Morris water maze test. n = 9 mice for WT and 10 mice for KO, ns, not significant, repeated measures two-way ANOVA and Student’s t-test and Mann-Whitney U test. (I–K) Normal fear learning and memory in *Lrfn2*^*-/-*^ mice (4–5 months). Mice with acquired fear memory in a spatial context combined with a sound cue (I) were tested for contextual fear memory 24 hours after training (J) and for cued fear memory 28 hours after training (K). n = 14 mice for WT and 17 mice for KO, ns, not significant, repeated measures two-way ANOVA and Student’s t-test and Mann-Whitney U test.

### Enhanced acoustic startle, but normal pre-pulse inhibition and susceptibility to induced seizure, in *Lrfn2*^*-/-*^ mice

Finally, to assess behaviors in sensory and motor domains, we first measured acoustic startle responses. *Lrfn2*^*-/-*^ mice showed enhanced startle responses to stimuli in a high loudness range (>110 dB) (**Figure 11A**). In contrast, pre-pulse inhibition was normal in *Lrfn2*^*-/-*^ mice, despite a tendency toward a decrease (**Figure 11B**), suggesting that sensory motor integration is normal.

**Figure 11.**
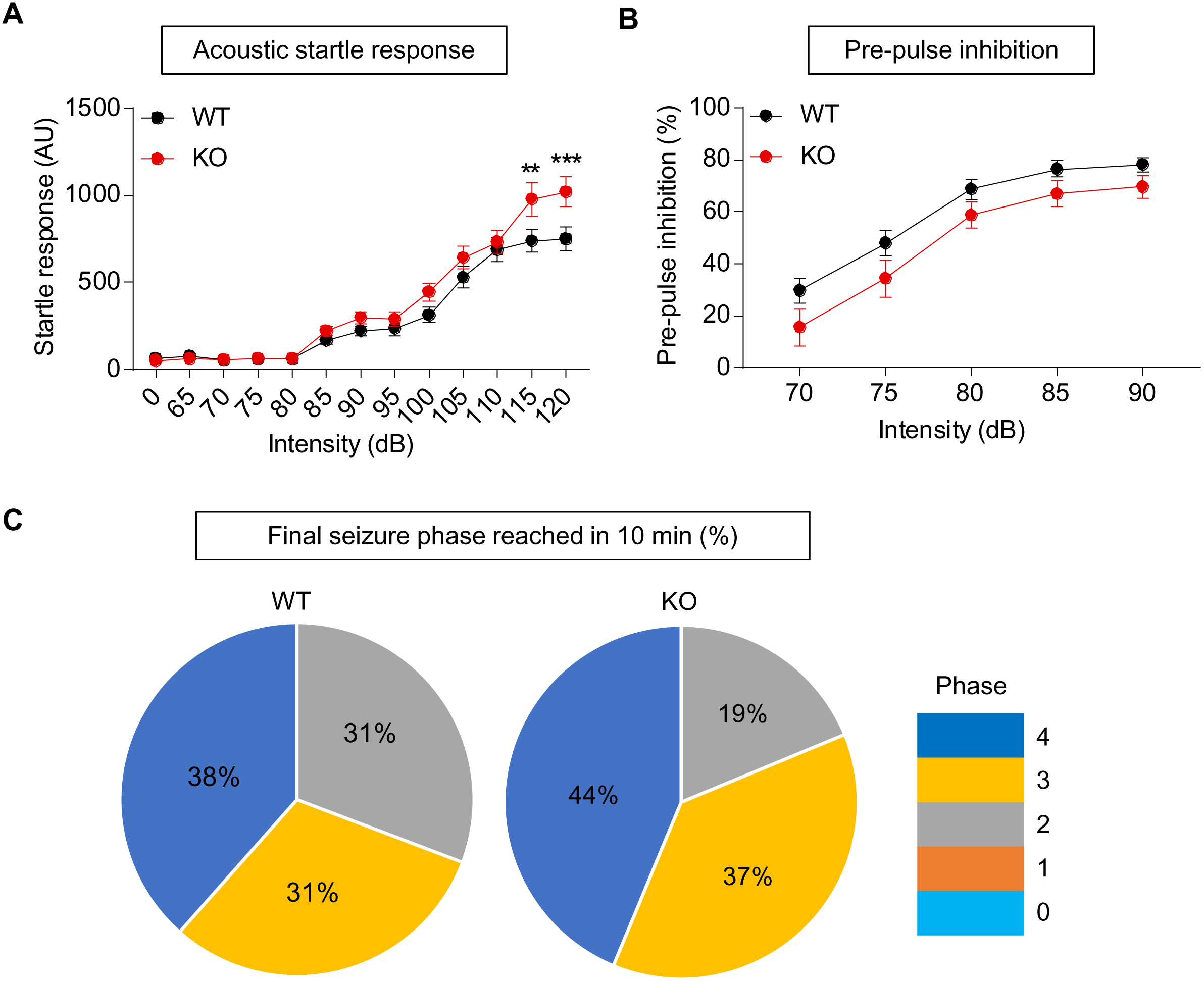
Enhanced acoustic startle, but normal pre-pulse inhibition and susceptibility to induced seizure, in *Lrfn2*^*-/-*^ mice. (A) Enhanced acoustic startle responses of *Lrfn2*^*-/-*^ mice (4–5 months) in a high loudness range (> 110 dB). n = 20 mice for WT and 24 mice for KO, **p < 0.01, ***p < 0.001, repeated measures two-way ANOVA with Bonferroni test. (B) Normal pre-pulse inhibition in *Lrfn2*^*-/-*^ mice (4–5 months). n = 14 mice for WT and 17 mice for KO, repeated measures two-way ANOVA with Bonferroni test. (C) Normal susceptibility to a seizure induced by PTZ (40 mg/kg; i.p.) in *Lrfn2*^*-/-*^ mice (5–6 months). The numbers in pie charts indicate the proportions of mice that reached the indicated stages of seizure at the end of the test (10 min). n = 13 mice for WT and 16 mice for KO, Chi-square analysis.

In a test measuring seizure susceptibility, *Lrfn2*^*-/-*^ mice showed normal susceptibility to seizures induced by pentylenetetrazolium (PTZ) compared with WT mice (**Figure 11C**). These results collectively suggest that a SALM1 deficiency leads to sensory hypersensitivity, but has minimal impact on sensory motor integration or seizure susceptibility.

## Discussion

### A SALM1 deficiency alters NMDAR-dependent synaptic plasticity

SALM1 has been shown to form a complex with NMDARs in vitro and in vivo and regulate dendritic surface clustering of NMDARs in cultured hippocampal neurons (Wang et al., 2006). We thus expected that a SALM1 deficiency in mice would lead to a reduction in NMDAR function. However, our results indicated that a SALM1 deficiency causes an increase in NMDAR function in the hippocampus, as evidenced by an increase in NMDA/AMPA ratio and normal basal transmission (a measure of AMPAR-dependent function).

This unexpected increase in NMDAR function led us to predict enhanced NMDAR-dependent synaptic plasticity in *Lrfn2*^*-/-*^ mice. However, we found that, although TBS-LTP was normal, NMDAR-dependent HFS-LTP and LFS-LTD were significantly reduced. What might explain these results? One possibility is that abnormally enhanced NMDAR-mediated synaptic transmission at *Lrfn2*^*-/-*^ excitatory synapses caused secondary changes in the molecular pathways downstream of NMDAR activation, suppressing NMDAR activation-dependent recruitment or removal of AMPARs during synaptic plasticity. Alternatively, the increase in NMDAR function may merely represent a compensatory change induced by insufficient synaptic delivery or removal of AMPARs during synaptic plasticity. These possibilities, which are not mutually exclusive, would create a situation in which basal NMDAR function is enhanced and activity-dependent NMDAR function during synaptic plasticity is suppressed. Given that SALM1 associates with NMDARs in vivo and promotes dendritic clustering of NMDARs in cultured neurons (Wang et al., 2006), it is unlikely that SALM1 deletion would markedly increase basal NMDAR function, making the latter possibility more likely. Although further details remain to be determined, our in vivo results clearly point to the possibility that SALM1 is required for bidirectional activity-dependent changes in AMPAR-mediated synaptic transmission during NMDAR-dependent synaptic plasticity.

In contrast to these significant changes in NMDAR-dependent synaptic plasticity, multiple lines of evidence suggest that a SALM1 deficiency has little effect on the development of excitatory synapses, at least in the hippocampus. First, the density of excitatory synapses was normal in the *Lrfn2*^*-/-*^ hippocampus, as supported by the normal density of excitatory synapses in EM analyses and the normal frequencies of mEPSCs and sEPSCs. In addition, the size and shape of excitatory synapses were minimally affected, as supported by the normal length, thickness, and perforation of PSDs in EM analyses, and the normal amplitudes of mEPSCs and sEPSCs. Therefore, an *Lrfn2* deficiency appears to have a greater effect on excitatory synaptic plasticity than on excitatory synapse development.

Notably, the findings reported here regarding excitatory synapses differ somewhat from those observed in the hippocampus of another *Lrfn2*^*-/-*^ mouse line (Morimura et al., 2017). The authors of this latter study reported major abnormalities in the morphology of excitatory synapses and dendritic spines in the hippocampus, with longer and thinner spines and more frequent oddly shaped spinule-like structures (Morimura et al., 2017). Our data, however, indicate no changes in the morphology of excitatory synapses, again as supported by the normal length, thickness, and perforation of the PSD, results that are in line with the normal amplitude of mEPSCs.

In terms of NMDAR function, this latter study found increases in the NMDA/AMPA ratio similar to those found here, but reported that LTP, an NMDAR-dependent function, was changed in the opposite direction: whereas our study found suppressed HFS-LTP, this previous study reported enhanced HFS-LTP. This reported increase in LTP could be attributable to the increased number of silent synapses observed in their mice, because silent synapses would have more room to accommodate incoming AMPARs during LTP. Our *Lrfn2*^*-/-*^ mice, however, appeared to have normal silent synapses, as supported by the normal frequency of mEPSCs. Therefore, silent synapses could not explain our LTP phenotype; moreover, LTP was suppressed rather than enhanced in our mice. However, it should be noted that NMDAR and LTP phenotypes in the two mouse lines arguably involve a similar defect, namely limited NMDAR-dependent synaptic delivery of AMPARs, although these defects appear to occur at different stages of synapse development: an early un-silencing stage of excitatory synapses and a relatively late post–un-silencing stage of excitatory synapse maturation.

Notably, both studies used mice that lacked the same exon 2 of the *Lrfn2* gene and had the same genetic background (C57BL6/J). Possible explanations for discrepancies in synapse phenotypes could include differences in conditions in which mice were bred and handled, the ages of mice used for slice recordings (P21– 35 in our study, and 3–6 months in the prior study), the method for preparing brain slices, or electrophysiology conditions (e.g., buffer solutions).

### A SALM1 deficiency suppresses inhibitory synapse development

An unexpected finding of our study was that an *Lrfn2* deficiency leads to suppression of inhibitory synapse development in the hippocampus, as supported by the decreased density of inhibitory synapses in EM analyses and the decreased frequency of mIPSCs and sIPSCs (Figures 4 and 5). It is possible that SALM1 could be targeted to inhibitory postsynaptic sites in CA1 pyramidal neurons, in addition to excitatory synapses, where it may regulate inhibitory synapse development, perhaps by participating in trans-synaptic adhesion with as yet unknown presynaptic ligands. Intriguingly, an EM analysis has shown that an *Lrfn2* deficiency causes widening of the synaptic cleft at excitatory synapses (Morimura et al., 2017).

Alternatively, SALM1 may be expressed in presynaptic GABAergic neurons. Indeed, our fluorescence in situ hybridization experiments revealed the presence of SALM1/Lrfn2 mRNA signals in cortical and hippocampal GABAergic neurons in addition to glutamatergic neurons. In GABAergic neurons, SALM1 may be targeted to presynaptic axon terminals, where it could participate in trans-synaptic adhesion or regulate presynaptic differentiation. In line with this possibility, SALM1 immunogold EM signals have been detected in presynaptic nerve terminals (Thevenon et al., 2016). Another possibility is that SALM1 in GABAergic neurons may be targeted to dendritic excitatory synapses. A lack of SALM1 at these sites might suppress excitatory synapse development or function, consequently suppressing output functions of GABAergic neurons.

### A SALM1 deficiency alters social communication and startle behavior

In our study, we found that *Lrfn2* knockout led to suppression of USVs in pups and enhancement of acoustic startle responses in adults. Our *Lrfn2*^*-/-*^ mice, however, were largely normal in other behavioral domains, including locomotion, anxiety-like behavior, social interaction, repetitive behavior, learning and memory, and seizure propensity. Behavioral test results reported for the previous *Lrfn2*^*-/-*^ mouse line (Morimura et al., 2017) show some overlap with ours, but are largely dissimilar. The most similar reported behavior is enhanced acoustic startle; the concurrence of the two studies on this point strongly suggests that SALM1 is required for normal development of auditory startle responses. The prior study did not observe any changes in pup USVs, another strong phenotype observed in our mice. Behaviors that were uniquely observed in the previously reported *Lrfn2*^*-/-*^ mouse line include suppressed social interaction, enhanced repetitive behavior, and enhanced learning and memory (Morimura et al., 2017). Again, these discrepancies could be attributable to differences in experimental conditions, including mouse breeding and handling. Importantly, the time at which behavioral experiments were performed differed between the two studies: light-off period for our study, and light-on period for the prior study.

The acoustic startle response has been shown to involve oligo-synaptic circuits and a small group of giant neurons in the caudal nucleus of the pontine reticular formation (PnC) (Yeomans and Frankland, 1995; Koch and Schnitzler, 1997), a brain region where SALM1/Lrfn2 mRNA signals are detectable, albeit at relatively low levels; moreover, current images lack sufficient cellular resolution to provide reliable information on this point (Figure 3 and Allen Brain Atlas). In addition, inhibition of NMDARs in the PnC by local infusion of APV has been shown to markedly suppress the acoustic startle response in rats (Miserendino and Davis, 1993). Given the abnormally enhanced NMDAR function in the *Lrfn2*^*-/-*^ hippocampus, it is tempting to speculate that SALM1 deletion in the PnC might lead to abnormal increases in NMDAR function and acoustic startle response.

USVs are an important mode of social communication in rodents, and deficits in USV have been observed in many mouse models of ASD (Scattoni et al., 2009; Wohr, 2014). Intriguingly, our *Lrfn2*^*-/-*^ mice showed suppressed isolation-induced USVs in pups, but normal female-induced USVs in adults. This difference may be attributable to the distinct nature of the two USV types: pup USVs are more anxiety‐ and development-related behaviors, whereas adult USVs are more reproduction related (Scattoni et al., 2009). Indeed distinct pup and adult USV phenotypes have been reported in neuroligin-2–knockout mice, which also show suppressed pup USVs but normal adult USVs (Wohr et al., 2013).

Pup USVs are known to involve many brain regions, including the periaqueductal gray (PAG) and amygdala (Hofer, 1996), where SALM1/Lrfn2 mRNA is expressed (Figure 3 and Allen Brain Atlas). In addition, NMDAR inhibition has been shown to suppress isolation-induced pup USVs in rats and mice (Winslow et al., 1990). Mouse pups also respond similarly, although certain NMDAR antagonists seem to induce biphasic responses; enhanced and suppressed USVs at low and high concentrations, respectively (Takahashi et al., 2009). Therefore, SALM1 deletion in mice might abnormally enhance pup USVs through NMDAR hyperactivity in brain regions that include the PAG and amygdala. In addition, because SALM1/LRFN2 has been implicated in ASD (Morimura et al., 2017), the reduction in pup USVs in *Lrfn2*^*-/-*^ mice may represent a novel autistic-like behavior. This is a potentially valuable tool because early symptoms are relatively common in human ASDs, but early behavioral measures are rare in animal models of ASD (Silverman et al., 2010; Wohr and Scattoni, 2013).

In conclusion, our results suggest that SALM1 is important for NMDAR function at excitatory synapses, and is required for normal synapse development at inhibitory synapses. Behaviorally, SALM1 is required for normal pup USVs and acoustic startle, but not for other behaviors tested.

## Acknowledgments

This work was supported by the National Research Foundation of Korea (NRF), funded by the Ministry of Science, ICT & Future Planning (NRF-2017M3C7A1048566 to H.K.), the NRF Global PhD Fellowship Program (NRF-2015H1A2A1033937 to R.K), the NRF (NRF-2017R1A5A2015391 to Y.C.B. and 2016R1A2B200682 to J.K.), and the Institute for Basic Science (IBS-R002-D1 to E.K.).

